# Temporal analysis of relative distances (TARDIS) is a robust, parameter-free alternative to single-particle tracking

**DOI:** 10.1101/2023.06.07.544011

**Authors:** Koen J.A. Martens, Bartosz Turkowyd, Johannes Hohlbein, Ulrike Endesfelder

**Affiliations:** Institute for Microbiology and Biotechnology, Rheinische-Friedrich-Wilhelms-Universität Bonn, Meckenheimer Allee 168, 53115 Bonn, Germany; Department of Physics, Carnegie Mellon University, 5000 Forbes Avenue, Pittsburgh, PA 15213, United States; Laboratory of Biophysics, Wageningen University and Research, Stippeneng 4, 6708 WE Wageningen, The Netherlands; Microspectroscopy Research Facility, Wageningen University and Research, Stippeneng 4, 6708 WE Wageningen, The Netherlands

## Abstract

In single-particle tracking (spt), individual particles are localized and tracked over time to probe their diffusion and molecular interactions. Temporal crossing of trajectories, blinking particles, and false-positive localizations present computational challenges that have remained difficult to overcome. Here, we introduce a robust, parameter-free alternative to spt: TARDIS. In TARDIS, an all-to-all distance analysis between localizations is performed with increasing temporal shifts. These pairwise distances fall in two categories: inter-particle distances not originating from the same particle, and intra-particle distances originating from the same particle. Since the distribution of inter-particle distances is unaffected by temporal shifts, the distribution of particle jump distances can be analytically fitted by analysing multiple temporal shifts. TARDIS outperforms existing tracking algorithms especially in complex conditions, and is even robust in scenarios that exceed the capabilities of current localization algorithms. Using TARDIS, we further show that measurements can be five-fold shorter without loss of information.

## Introduction

Single-particle tracking (spt) is a powerful technique in which individual particles are followed through time to infer information about a conjugated particle of interest, or about the environment in which the particles are embedded^1–3^. By attaching a contrast agent to a particle of interest, spt can be used to assess molecular dynamics *in vitro* or *in vivo*. Applications of spt include, but are not limited to, photo-activatable localization microscopy^4^ (sptPALM) via photo-activatable organic fluorophore or fluorescent protein conjugation, or interferometric scattering microscopy (iSCAT) via e.g. gold nanoparticle conjugation^5,6^. These methods enable spt with < 40 nm spatial and > 100 Hz temporal accuracy. However, especially endogenous *in vivo* spt is limited to using sptPALM with fluorescent proteins, which limits the trajectory length to around 3-10 timepoints^4,7^.

Analysis of spt(PALM) is generally divided in two steps: (1) detection of individual particles and localization with sub-pixel accuracy, and (2) temporal linking of these localizations to trajectories. Detection and superresolved localization (step 1) has seen big improvements in the recent decade following breakthroughs in computational image analysis^8–11^, machine learning^12–14^, and microscopy hardware design^15–18^. Accurate static particle localization at high densities and low signal-to-noise has become feasible and performance of localization algorithms as well as the required computational hardware will likely continue to increase further.

The second analysis step (tracking), however, is impeded by high computational complexity, especially when considering global information and taking possible state-changing, blinking, and bleaching behaviour into account^19,20^. Currently, most spt(PALM) experiments are designed to reduce this computational complexity by using a very low density of non-blinking particles (i.e., <0.1 per μm^2^) over long data acquisition times^2,19^. Even with these considerations, tracking algorithms require *a priori* parameters, such as the likelihood of state-changing or bleaching, maximum search radius, and expected blinking behaviour; still, the obtained trajectory information could unknowingly contain imperfect linkages. The resulting trajectories can additionally be deteriorated by spurious localizations, which is often unavoidable in livecell imaging^21^.

The trajectories are then quantitatively interpreted by, e.g., analysing jumping distance (JD) histograms, mean squared displacement (MSD) curves, or by performing diffusion distribution analysis (DDA)^7,22,23^. However, not all of these interpretations fully utilize the information within a trajectory – for instance, a JD histogram analysis does not require information on the complete trajectory, only whether or not two localizations are linked together on consecutive frames. Taken together, tracking algorithms do not take full advantage of recent developments in (high-density) particle localization, have high computational complexity, can unknowingly introduce linkage errors, and full trajectory information is not always utilized to obtain state-of-the-art biological knowledge.

Here, we propose an analysis method that does not require *a priori* assumptions, cannot introduce linkage errors, and is minimally influenced by localization density, particle blinking, and random or structured spurious localizations. Our method to analyse tracking data will open the way for new experimental avenues by allowing higher particle densities, a wider pool of possible particles, and lower impact of spurious background localizations. We focus on fluorescence sptPALM data throughout this manuscript, but note that TARDIS is method-agnostic and can be applied to all spt data.

TARDIS (temporal analysis of relative distances) performs a global analysis on all relative spatiotemporal distances, inspired by methods that infer localization precision or structures from relative distances of particles^24–26^. TARDIS does not use any tracking algorithm and thus circumvents required inherit assumptions and experimental considerations. We show that TARDIS provides accurate quantitative measures and outperforms existing tracking algorithms in complex spt conditions, such as high-density localization, heavily noise-deteriorated situations, blinking particles, and undergoing intricate, non-Brownian, diffusive motion. TARDIS can accurately deduce unknown jump distance distributions, or analytically described single and double Brownian-motion populations, state-switching populations, and can be expanded to other analytically-described conditions. We provide TARDIS as a ready-to-use software, both as stand-alone GUI and as a MATLAB function for incorporation in analysis routines (https://github.com/kjamartens/TARDIS-public).

## Materials and methods

### Simulation of single and dual diffusive species

A redundant number of trajectories with an underlying specified diffusion coefficient and 30 nm localization precision (ignoring effects from motion blur) were simulated in a two-dimensional plane. The trajectories had a length determined via a three-frame half-life, with a frame-time of 10 ms, on a field of view of 10 x 10 μm. To simulate blinking particles (where indicated), localizations were randomly removed from these trajectories. This localization list was then limited to the number of localizations that corresponds to 10.000 un-filtered trajectories. The trajectories were given a random starting frame, where the number of frames was varied to create a certain localization density. Then, spurious localizations were added, the amount of which were based on the specified values and the number of localizations in the trajectories. These spurious localizations were randomly spread throughout the field of view and frames. For testing TARDIS with two diffusive populations, the same procedure was used, but all trajectories were randomly sampled from one of two populations with corresponding diffusion coefficients.

TARDIS was run on these datasets with the following settings (see Supplementary Note 1 for a full explanation of these parameters): maximum jump distance 5 μm, max τ 3 frames, 10 frames longest trajectory length (determined via the Wilcoxon test), integration of 50 frames used for higher background subtraction accuracy, 30 nm localization precision, diffusion coefficient start value random between 0.1 and 10 μm^2^/s, using an estimation fit.

### Simulation of state-switching species

For all specified conditions in the state-switching species (spurious and membrane localizations in 1:0, 1:0.5 and 1:1 ratio with respect to true localizations), ten repetitions of a simulation similar to ones described previously with the software encompassed in anaDDA^7,22,23^ were created with the following settings: 40 (bacterial) cells (simulated as cylinders with length random between 2-3 μm and radius random between 0.45 and 0.55 μm, capped by two half-spheres with a radius identical to the cylinder radius; the cells were rotated and translated randomly throughout a 25 x 25 μm FoV) were populated with [300-500] particles in 2000 frames (10 ms frame time). Trajectory lengths were pulled randomly from a decay with half-life = 3 frames (limited at 8 frames). Localizations that were outside the FoV were discarded. Every particle was simulated with *k*_on_ = 50 s^-1^ or 20 s^-1^, *k*_off_ = 30 s^-1^ or 20 s^-1^, and *D*_free_ = 2 μm^2^/s, and localizations were subject to a 30 nm localization error. Simulation was performed with the software encompassed in anaDDA^22^, in which a 1e-8 s steptime and a precision factor of 50000 was used. Starting frames for trajectories were randomly assigned. Spurious localizations were added where needed based on the number of true localizations, and positioned randomly throughout the FoV. Membrane localizations were added where needed based on the number of true localizations, and positioned in a 2-dimensional, 100 nm diameter slice at the simulated cell outlines.

All datasets were analysed with TARDIS using the following settings (see Supplementary Note 1 for a full explanation of these parameters): anaDDA fit model with 0.5 μm radius and 3 μm length, 10 ms frame time, 30 nm localization precision. Maximum jump distance calculated was 2.5 μm (i.e., average cell length), max Δt was 5 frames, 87 frames for longest trajectory length (determined via the Wilcoxon test) using the integration of 50 frames for higher background subtraction accuracy. anaDDA starting parameters were randomly chosen from [10-80] s^-1^, [10-80] s^-1^, [1-4] μm^2^/s, and an estimation fit (i.e, a fit on remaining JDs after subtraction of inter-particle fraction; see Supplementary Note 1) was performed.

### Spt(PALM) microscopy

All microscopy was performed on home-build microscopes as described in detail elsewhere^27^ that allows for imaging of Dendra2 proteins via primed conversion^28^. Briefly, live cell experiments were performed on the following system: 561 nm (OBIS, Coherent Inc., USA), 488 nm (Sapphire LP, Coherent Inc., USA), and 730 nm (OBIS, Coherent Inc., USA) were combined and controlled via an acousto-optical tuneable filter (except the 730 nm laser) (AOTF; TF525-250-6-3-GH18, Gooch and Housego, USA). The lasers were directed towards the back-port of a Nikon Ti Eclipse microscopy body, and focused on the back-focal plane of a CFI Apo TIRF 100x objective (NA 1.49, Nikon) in HiLo mode via an ET dapi/Fitc/cy3 dichroic), and emission light passed through a ZT 405/488/561rpc rejection filter and ET610/75 bandpass (all AHF Analysentechnik, Germany). The emission light was focused on an iXON Ultra 888 EMCCD camera (Andor, UK) with an effective pixel size of 128 nm. The image acquisition was controlled by micromanager^29^.

Bead experiments were performed on a similar microscope setup: A 638 nm laser line (Novanta, Boston, MA, USA) was controlled via a TriggerScope 4 (Advanced Research Consulting, Newcastle, CA, USA) connected to PycroManager^30^, and directed towards the back-port of the Nikon Ti body via a laser-coupled fibre (150 μm core diameter, Thorlabs). The laser light was directed via a reflective collimator and via a ZT405/488/561/642rpc and ZET405/488/561/642m-TRF dichroic/rejection filter combination (Chroma, Bellows Falls, VT, USA) to the back-focal plane of a 60x Apochromat TIRF 1.49 NA objective (Nikon) in HiLo mode. Emission light was passed through a 655 nm-longpass filter (655 LP ET, AHF Analysentechnik, Germany) and collected on a Prime BSI sCMOS camera (Teledyne Photometrics, Tucson, AZ, USA; 107 nm effective pixel size).

For the bead diffusive experiments, Carboxylate-modified FluoroSpheres (715/755 excitation/emission peaks, 97 nm diameter according to manufacturer) (Invitrogen, Thermo Fisher), were diluted 2500 times or 10000 times in purified water. After 5 minutes sonication, these solutions were placed in an Ibidi 8-chamber-well slide (Ibidi, Gräfelfing, Germany), and imaging was performed in a temperature-controlled (20 ± 0.5 °C) room with the optical settings described above.

For the live-cell experiments, *E. coli* MG1655 harbouring a rpoC-Dendra2-strain^31^ was inoculated into fresh LB medium and incubated overnight at 37°C while shaking (180 RPM). On the day of the experiment, cells were reinoculated into fresh EZ Rich Defined Media (EZRDM, Teknova) and incubated at 37°C while shaking (180 RPM) for one hour. Next, the culture was split into two subcultures, and to one of them, rifampicin was added to a final concentration of 300 μg/mL. Both cultures were incubated at similar conditions for one hour and centrifuged for 2 min 3000 x g afterward. The supernatant was discarded, cell pellets were washed with fresh EZRDM and centrifuged again with similar parameters and the supernatant was discarded. Cell pellets were resuspended in the residual supernatant and 2 μL of cell suspension was placed on 1% EZRDM-agarose pads (see below) for imaging.

Low melting agarose (Merck-Sigma Aldrich) was suspended in fresh EZRDM to a final concentration of 1% and incubated for 12 minutes at 70°C until the solution was clear, then the agarose solution was stored at 42°C for further use. To prepare agarose pads, 100 μL of agarose solution was placed on an indented microscope slide and covered with a coverslip (#1.5, Marienfield), which was prior cleaned overnight with 1 M KOH (Carl Roth) solution, and incubated for 2 hours at room temperature. Then, the coverslip was discarded, and the cell suspension was loaded on the solid agarose pad and covered with a new, clean coverslip.

## Results

### The TARDIS algorithm

Our proposed temporal analysis of relative distances (TARDIS) method is a parameter-free algorithm that only requires spatiotemporal information of particle localizations (Figure 1a). TARDIS connects all localizations on frame N to all localizations on frame N+τ, where the temporal shift τ is gradually increased (Figure 1a,b). TARDIS then records the jump distances, visualised here as a histogram. At small values of τ (i.e., τ smaller or equal to the observed trajectory length of a particle), the linkages between localizations fall in two distinct categories: 1) linkages in which both localizations belong to the same particle (blue lines in Figure 1b; intra-particle links) or 2) linkages in which the localizations do not belong to the same particle (red lines in Figure 1b; inter-particle links). Importantly, at large values of τ, i.e., τ larger than the maximum trajectory length in the dataset, only inter-particle links (red) will be obtained (Figure 1b, right). The underlying inter-particle links distribution is insensitive to time, i.e., it is does not vary over values of τ. Therefore, the intra-particle links contained in the data obtained at small τ (Figure 1b, bottom) can be extracted. This can be done by subtracting the inter-particle links distribution found at large τ (red curve) (called ‘TARDIS-JD’ from here on), or, ideally, it can be fitted with a combination of the inter-particle links distribution along with an analytical distribution describing the intra-particle links (blue curves). This combined fitting routine is less sensitive to noise and provides more accurate fitting results compared to TARDIS-JD. Starting parameters for the full TARDIS routine can be estimated by first performing TARDIS-JD (‘estimation fit’). Alternatively, TARDIS-JD extraction can be used independently to obtain JD distributions for datasets that contain dynamics that have not yet been analytically described, such as complex multi-state dynamics and/or directed motion, or obtained from unknown underlying dynamics. An in-depth description of the TARDIS software implementation is given in Supplementary Note 1.

**Figure 1:**
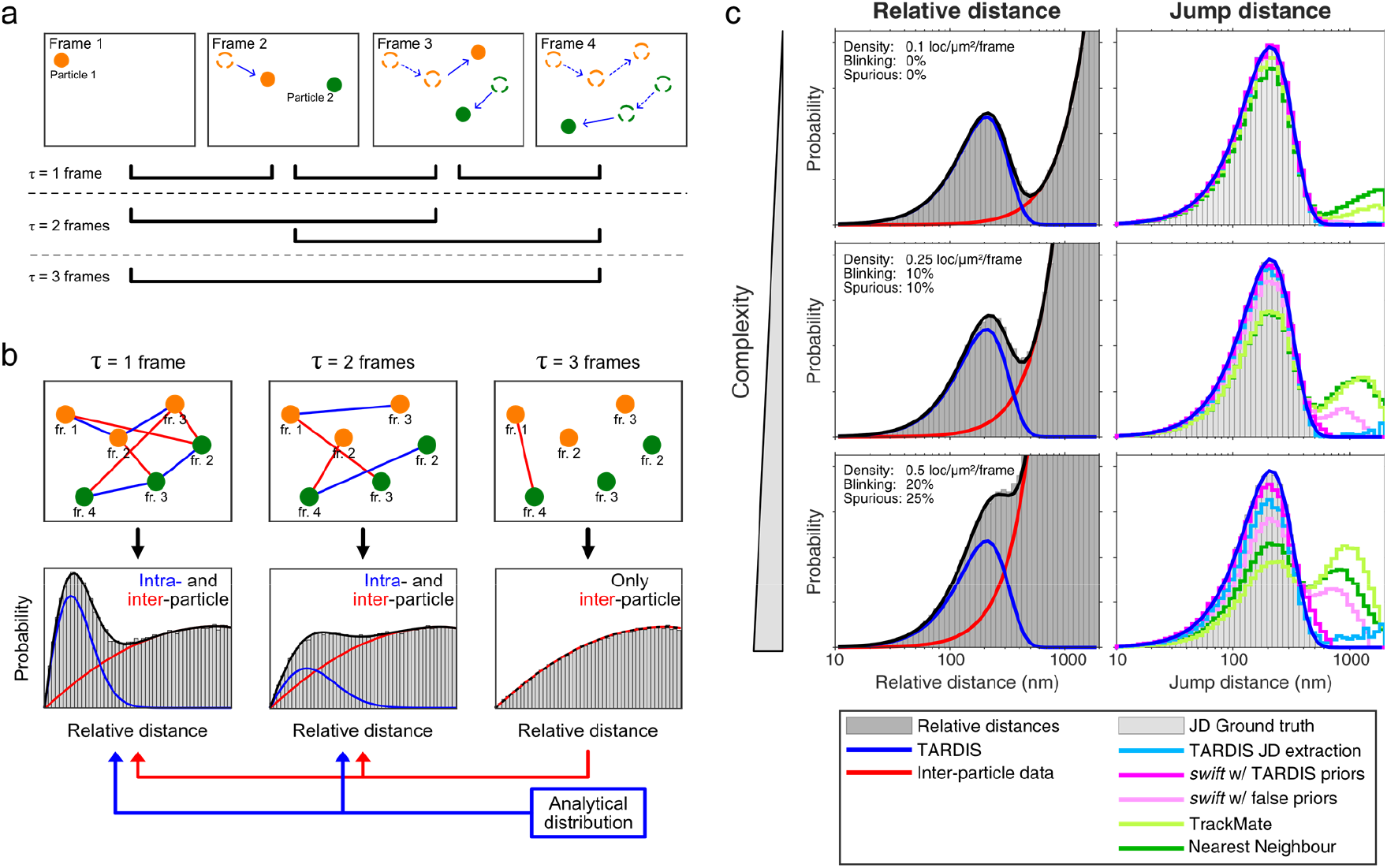
Overview of the TARDIS algorithm. (**a**) To exemplify the TARDIS algorithm, two diffusing particles are simulated, starting on frame 1 and 2, respectively. The spatiotemporal positions of the localizations are recorded. TARDIS compares the positions on all frames with a set time-shift τ, here illustrated for τ = 1, 2, and 3. (**b**) All localizations at time t are compared to all localizations at multiple times t+τ over the complete temporal range, illustrated for τ = 1, 2, and 3. At τ smaller than the longest trajectory (i.e. τ < 3 in this example), the distribution that can be determined from the relative distances contains information from both intra- and inter-particle links (blue and red lines, respectively. Using a temporal shift that is larger than the longest trajectory (i.e., τ ≥ 3 in this simulated example), only inter-particle links are present in the distribution (red lines). This inter-particle links distribution is used in conjunction with an analytical distribution (a single population fit is shown here) to fit the distributions obtained from temporally analysed relative distances. Distributions in b are created via a simulation of 25,000 particles. (**c**) Comparison of TARDIS with single-particle tracking algorithms. Three simulated (20.000 trajectories) datasets (10 ms frametime) with increasing complexity (by increasing localization density, particle blinking, and spurious localization density) are analysed with TARDIS (left; τ = 1 frame shown here) and compared (right) with *swift* (Endesfelder et al., manuscript in prep.) (magenta), TrackMate^32,33^ (light green) and Nearest neighbour analysis (dark green).

Any analytical distribution which describes jump distances over multiple τ values can generally be used in TARDIS. We employ a single-population, freely-diffusive fit in Figure 1. Additionally, the rate at which the intra-particle links fraction decreases as a function of τ can be used as a measure for bleaching kinetics (Supplementary Note 2). Further, the value of τ at which only inter-particles are present can be estimated without *a priori* information via a one-sided Wilcoxon statistical test (which deduces the value of τ at which the JD distributions do no longer change as a function of τ; Supplementary Note 3), which makes TARDIS a spt analysis method that does not require any *a priori* parameters. Finally, TARDIS can provide the user with statistical estimates about particle parameters, such as mean jump distance, blinking probability, and average trajectory length (Supplementary Note 4) The TARDIS software is available as open-source MATLAB code and as a stand-alone GUI program (https://github.com/kjamartens/TARDIS-public).

### TARDIS outperforms existing single-particle tracking methods

Current state-of-the-art static localization algorithms can localize immobile particles with up to ∼5 localizations/μm^2^/frame densities^13,14^, whereas current tracking algorithms normally operate at << 0.1 localizations/μm^2^/frame, indicating a big gap in accessible densities. Further factors complicating tracking algorithms are the blinking of (fluorescent) particles and the presence of localizations not belonging to trajectories. These can either arise from experimental properties (e.g., photo-physics, autofluorescence), but also from faulty localization, especially when combining blurry, mobile point spread functions with multi-particle fitting (Discussion). To investigate the performance of TARDIS in these complex spt(PALM) conditions, we simulated (Materials and methods) datasets of particles with increasing particle density, particle blinking, and spurious localization density, which all effectively increase the amount of inter-particle linkages relative to intra-particle linkages, hereafter called ‘complexity’ (Figure 1c, left, Supplementary Note 5).

The simulated datasets were analysed (Figure 1c, right, also see Extended Data figure 1) with TARDIS (blue, cyan) and compared to the jump distance (JD) ground truth (grey histogram), as well as to analysis with a range of tracking algorithms with a varying degree of inherit assumptions and *a priori* information: nearest-neighbour tracking analysis (dark green), which is robust at low complexity, but has a chance for wrong linkages at higher complexity; (2) Linear Assignment Problem (LAP) tracker embedded in TrackMate^32,33^ (light green), which performs a spatiotemporal global optimization protocol, and is especially optimized for Brownian motion; and (3) a global Bayesian tracking method (*swift* (Endesfelder et al., manuscript in prep., beta-testing repository http://bit.ly/swifttracking), magenta), which relies on providing *a priori* estimations for all parameters and iterates over these. It should be noted that TARDIS can provide estimations on tracking parameters to assist the complex tracking algorithms (Supplementary Note 4).

TARDIS shows excellent agreement with the ground-truth throughout the tested parameter space, also when only extracting the JDs (TARDIS JD extraction, cyan in Figure 1c, right) rather than fitting the population. *swift* (magenta) was performed with two different *a priori* information: either static, ‘false’ priors (light magenta) or dynamic priors determined via TARDIS (mean jump distance, bleach time, and blinking chance; see Supplementary Note 4). *swift* performs well when using meaningful *a priori* information, e.g., TARDIS-generated. Other tracking algorithms behave adequately at low complexity, but show non-existing high jump distance at increased complexity, due to incorrect inter-particle linkages, which can be attributed to inherit ‘greediness’ of these tracking software to link localizations together, rather than terminating trajectories. We also observed under-estimation of jump distance via the tracking algorithms at high densities and high diffusion coefficients, which is a result from linking localizations from different particles together (Extended Data figure 2).

### TARDIS is largely unaffected by inaccurate localization data and high particle densities

As the TARDIS algorithm performs well under complex scenarios, we investigated to what extend TARDIS remains accurate. We investigated (1) well-localized single-particle trajectories across a range of diffusion coefficients (with 10 ms frametime) and densities (Figure 2a,d), (2) the influence of spurious noise and blinking particles (i.e. ‘localization inaccuracies’) (Figure 2b), and (3) multiple diffusive populations combined with previously established complexity (Figure 2c).

**Figure 2:**
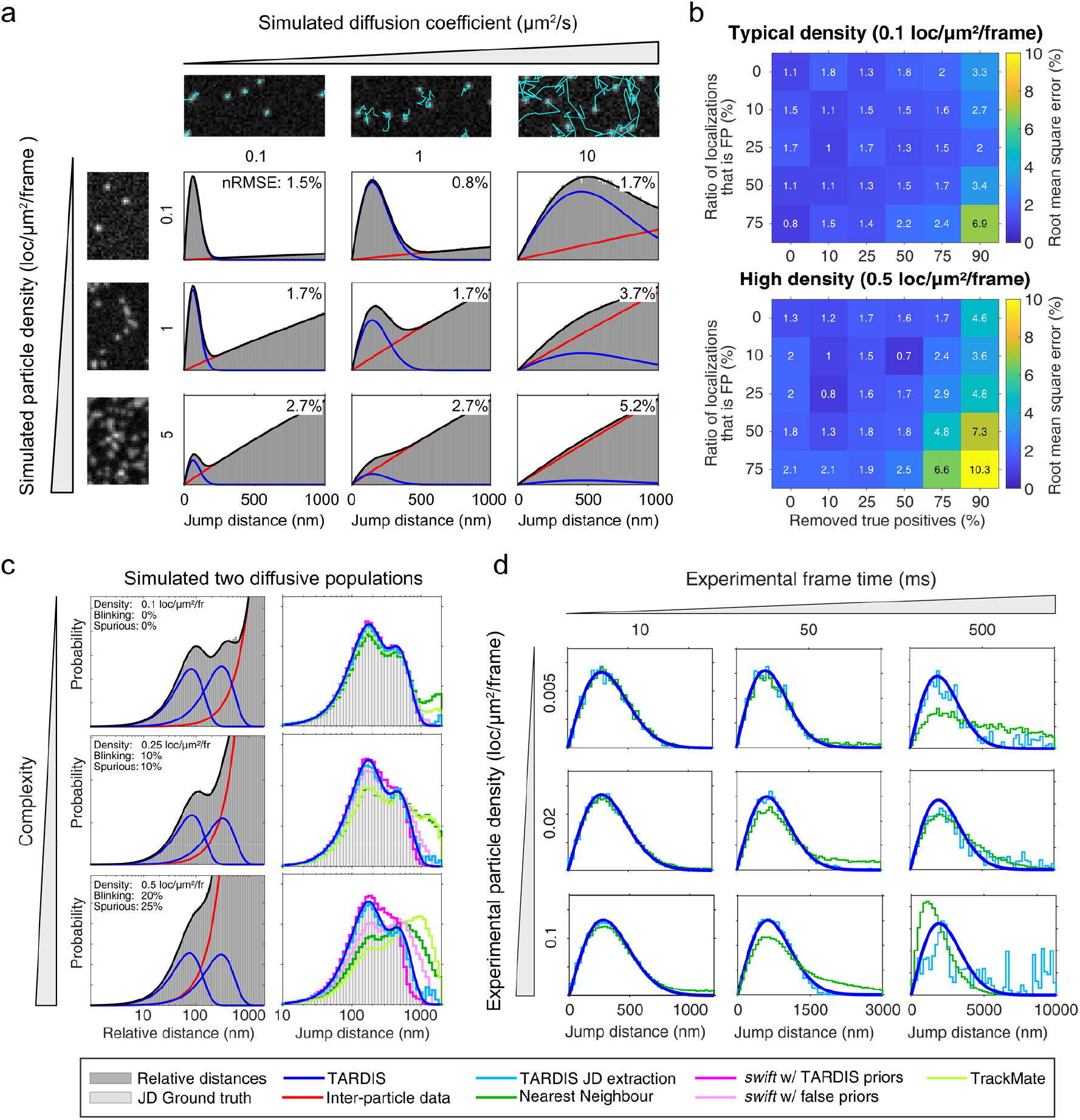
Performance of TARDIS in conditions of increasing complexity. (**a**) Visualisation of a TARDIS fit across a range of particle densities (vertical) and diffusion coefficients (horizontal). TARDIS is performed on 20.000 simulated localizations with 30 nm localization precision; images on the left show representative particle density, images on the top show representative jump distances (over 20 frames) in cyan. The simulations and TARDIS fits were repeated ten times, and the normalized root mean square error (nRMSE; normalized to diffusion coefficient) from these iterations is reported in the top-right of every sub-plot. Detailed information about the fits and comparison with tracking algorithms can be found in Extended Data figures 2 and 3. (**b**) Error of the TARDIS fit of a single diffusive population (with a bleaching half-time of three frames) at different particle densities and different levels of localization inaccuracy. Simulations of particle trajectories with a diffusion coefficient at 1 μm^2^/s at densities of 0.1 and 0.5 loc/μm^2^/frame were degraded by randomly removing true positive (TP) localizations from the trajectories and/or adding unstructured false positive (FP) localizations randomly throughout the field of view in various contributions. Reported values are calculated from 10 repetitions of every condition. Detailed information about the TARDIS fits can be found in Extended Data figures 3 and 4. (**c**) Fitting two populations in TARDIS scales similarly with complexity. Trajectories were simulated with, and analysed with, identical parameters as specified in Figure 1b, except half the trajectories were given a 0.5 μm^2^/s diffusion coefficient, and the other half a 5 μm^2^/s diffusion coefficient – and analysis with TARDIS was performed with two populations. The information is also presented on a linear x-axis in Extended Data figure 1. (**d**) Experimental single-particle mobility assessed by TARDIS compared to TARDIS-JD and nearest-neighbour tracking in a range of complexities by increasing the fluorophore concentration and increasing the frame time while keeping excitation time constant. To fully prevent multi-particle artefacts (see Supplementary Note 6), the localization list belonging to the highest density is created via the localizations of the lowest density. Throughout this parameter space, the same total experiment time was used (250 seconds).

We tested TARDIS’ accuracy by analysing simulated single-population diffusion with a varying degree of particle density and diffusion coefficient (Figure 2a). The complexity increases throughout the tested parameter space, as indicated by the increasing ratio of intra-particle and inter-particle links. TARDIS can accurately (< 5% nRMSE (normalized root mean square error)) compute single-population diffusion up to ∼50x higher particle densities than commonly used in spt(PALM) (5 localizations/μm^2^/frame), assuming localization is accurate at these high densities, with all tested varying diffusion coefficients (up to 10 μm^2^/s, localized every 10 ms) to further increase complexity (Figure 2a; for comparison with tracking methods, see Extended Data figure 2; detailed fitting results are presented in Extended Data figure 3). We note that the nRMSE of the found diffusion parameter scales qualitatively with the complexity, indicative that TARDIS will still be accurate at even higher complexity.

Incorporating inaccurate (i.e., false positive or false negative) localizations in our simulations (Figure 2b) reveals the insensitivity of TARDIS to both false positives (FP; e.g., spurious localizations) and missing true positives (TP; e.g., blinking particles, or particle moving temporarily out-of-focus). Diffusion coefficient (Extended Data figure 3) and bleach time (Extended Data figure 5) are accurately retrieved up to the condition where 50% of the localizations are removed (i.e., on average ∼1.5 localizations/particle or ∼1.2 consecutive localizations/particle) while simultaneously 50% of the dataset consists of FP localizations. In fact, a ∼10% (n)RMSE is still achieved when more than 99.9% of all information in the relative distance histograms is attributed to inter-particle linkages (Extended Data figure 4).

Next, we investigated the accuracy of TARDIS in conditions with complex datasets, which additionally have two diffusive populations rather than one (Figure 2c). TARDIS accurately fits the relative distance histogram with a mixture of inter-particle distribution and two diffusive populations, whereas the JD histogram obtained via tracking algorithms does not clearly show the separate populations. JD extraction in TARDIS (cyan curve in Figure 2c) does accurately show the JD ground-truth.

Finally, we assessed TARDIS under experimental conditions with increased complexity (Figure 2d, Extended Data figure 6). To do so, we imaged freely diffusing fluorescent ∼100 nm diameter beads in water, imaged with 200 μs stroboscopic illumination to minimize localization artefacts (Methods), while changing the bead density and frame time (i.e., increasing jump distance). These datasets were localized and analysed via TARDIS, TARDIS-JD extraction, and nearest-neighbour tracking. At more complex scenarios, nearest-neighbour tracking and TARDIS-JD fail to accurately deduce jump distances. Due to the global fitting nature of TARDIS (i.e., over multiple values of τ), the diffusion coefficient can still be accurately extracted (lower panels in Figure 2d). These results are very similar to the ones obtained via simulated trajectories.

Combining these results indicate that TARDIS is largely unaffected by regularly encountered particle, sample, or computational imperfections or influences that have a large impact on traditional spt(PALM) analysis accuracy. The scaling of TARDIS’ accuracy with complexity clearly indicates that experimental data will be hindered by localization artefacts sooner than by inaccurate TARDIS analysis.

### TARDIS accurately elucidates intricate diffusive behaviour

So far, we have focused on well-described single- or double-population Brownian motion. However, in biological conditions, this is normally not the case, and the diffusive behaviour is more intricate. Therefore, we assessed how TARDIS can handle diffusional complexity with three experiments (Figure 3).

**Figure 3:**
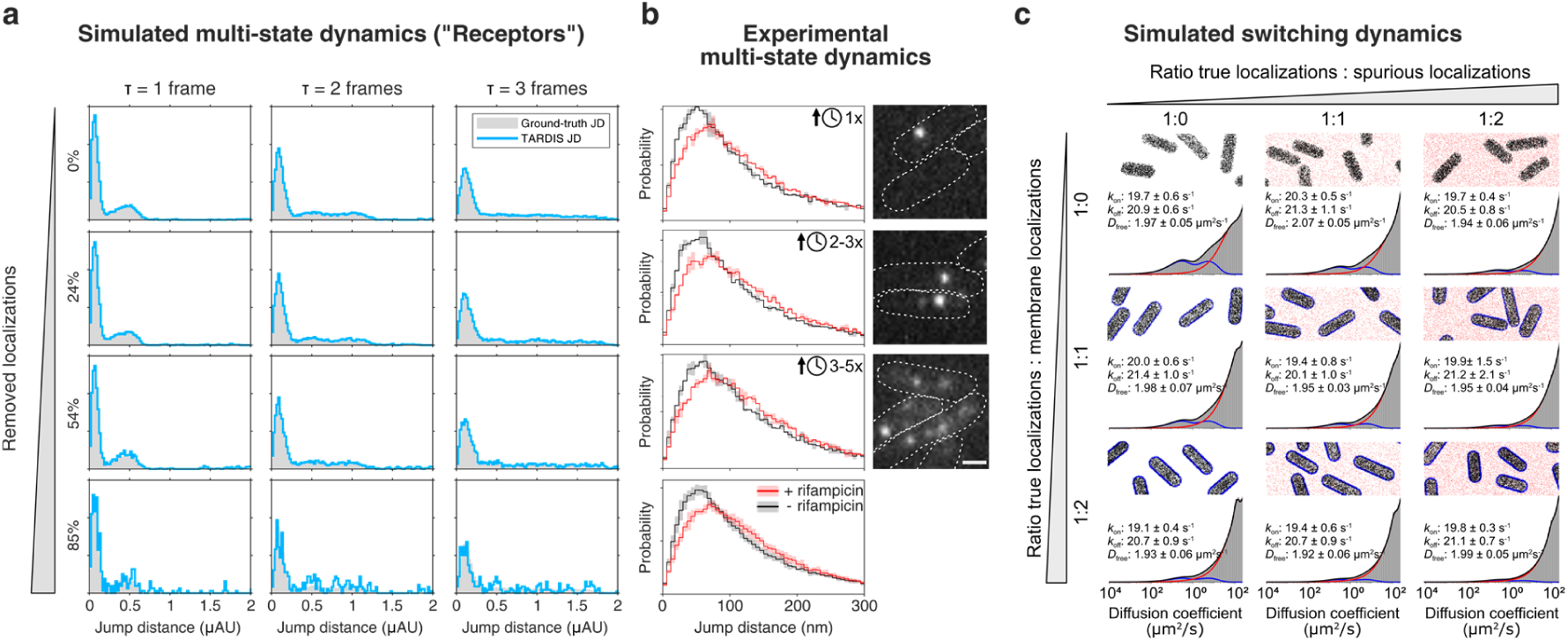
TARDIS enables high-complexity single-particle mobility experiments of intricate diffusive behaviour. (**a**) The “Receptors” tracking data from Chenouard et al^19^ (localization density set at 0.15 loc/μm^2^/frame) has been deteriorated by removing localizations to simulate lower signal-to-noise conditions similar to Chenouard et al^19^ (rows) and analysed via TARDIS JD extraction, and compared to the ground-truth (GT) data. The complex diffusive behaviour of these particles cannot be well-described analytically, as the JD histograms over time-shifts (columns) show. The following TARDIS settings were used: τ bins of 1-3 frames; maximum jump distance of 10 μAU. Other simulated datasets from this resource can be found in Extended Data figure 7. (**b**) Jump distance analysis of RNA polymerase in *E. coli* with (red) and without (black) rifampicin. By increasing the photo-conversion rate, the total time required to obtain 80 localizations per μm^2^ (within cells) is decreased by a factor of ∼2.5 (2^nd^ panel), then by a factor ∼4 (3^rd^ panel). Insets on the right show exemplary frames of these movies. The bottom panel combines the jump distance histograms of the top three panels. Lines indicate average of two biological replicates, shaded area indicates the standard deviation. Scalebar indicates 1 μm. (**c**) Fitting of a simulated state-changing population in bacterial cells (*k*_on_, *k*_off_ = 20 s^-1^, *D*_free_ = 2 μm^2^/s, three frame bleaching half-time, with trajectory length capped at eight frames). Inaccuracies from ground truth are introduced as spurious localizations (columns, red localizations) and membrane-bound localizations (rows, blue localizations). Reported values are median ± quartile determined from ten repetitions. Detailed information about the TARDIS fits can be found in Extended Data figure 9. Simulation datasets consist of 10.000 ‘true positive’ trajectories at 10 ms frame-time.

First, we analysed simulated datasets provided as part of a comparison of particle tracking methods^19^ (Figure 3a, Extended Data figure 7). Since the underlying biological model changes throughout the datasets, we used the TARDIS JD extraction. To investigate the effect of data complexity, we reshuffled the “Receptors” trajectory localizations (described as “tethered motion, switching, any direction”) to obtain a high density of particles (380 particles/frame or ∼0.15 loc/μm^2^). Additionally, we removed localizations according to the four signal-to-noise-ratio (SNR) levels (SNR = 7, 4, 3, and 1, equivalent to removing 0, 24, 54, and 85% of localizations, assuming perfect localization at SNR = 7, and scaling logarithmically) used in the comparison^19^. While the JD histograms show intricate behaviour over a range of temporal shifts, TARDIS JD extraction shows good agreement with the ground-truth for the SNR = 7, 4, and 3 levels (Figure 3a). The other simulated datasets from this resource were also analysed in this way (Extended Data figure 7), showcasing that TARDIS JD extraction can handle challenging datasets with intricate diffusive behaviour.

Next, we explored the use of TARDIS to drastically reduce imaging time in sptPALM. RNA polymerase in *E. coli* has dynamic diffusive behaviour that does not satisfy a known analytical model^34,35^. Rifampicin is a transcription inhibitor that decreases the promoter search and transcription kinetics, effectively increasing the diffusivity of RNAP^34,35^. Therefore, we used TARDIS-JD extraction and qualitatively assess the obtained jump distance histograms (Figure 3b). By increasing the primed conversion efficiency (i.e. increasing 488 nm laser power) in repetitions of the experiment^28,31^, we were able to reach the same total number of localizations per cell area (80 localizations per 1 μm^2^ cell area) much faster, decreasing the total experiment time step-wise ∼2-3-fold and ∼3-5-fold. Addition of rifampicin is characterised by a shift towards higher RNAP jump distances, which is clearly observed after TARDIS-JD extraction (Figure 3b). Even though the change in jump distance is only ∼30 nm, this small difference is consequently extracted via both low- and high-throughput sptPALM-TARDIS (Figure 3b, bottom panel), which is not the case when assessing this data via nearest-neighbour tracking (Extended Data figure 8). A further decrease of imaging time is hindered not by TARDIS, but rather by difficulties in the localization procedure, e.g., fitting overlapping, blurry PSFs (Discussion).

Finally, we investigated a scenario in which state-switching sptPALM in prokaryotic cells is simulated. The trajectories show transient binding with specific kinetics that can be extracted with diffusion distribution analysis (DDA) approaches^7,22,23^, after a translation from JD to apparent diffusion coefficient. The simulated data is deteriorated by spurious localizations on the cell membrane (i.e., structured noise), and additionally randomly throughout the field of view (i.e., unstructured noise) (Figure 3c, Extended Data figure 9). The ground-truth simulation parameters are recovered via TARDIS with small (< 5%) error with respect to using analytical DDA on the GT trajectories throughout the tested deterioration, even in the extreme case in which the ratio of [TP:spurious localizations:membrane localizations] is 1:2:2 (i.e., only 20% of localizations are GT trajectory information).

These experiments showcase that TARDIS enables measurements in experimental conditions that are currently largely inaccessible via tracking-based analysis, removing the current stringent metrics such as low-density, long-duration experiments, and non-blinking particles. TARDIS opens up the opportunity to obtain high-statistical diffusion information in shorter time, which in turn decreases the inherently present temporal averaging in single-particle experiments.

## Discussion

Single-particle tracking analysis is limited by both computational complexity and stringent experimental requirements. Reconstructing truthful particle trajectories from a dataset with spatiotemporally close trajectories is challenging without accurate *a priori* information, and is even further complicated by the blinking and/or fast bleaching of particles. TARDIS solves all these issues by analysing all-to-all particle distances over multiple temporal shifts. TARDIS accurately resolves single, double, and state-changing populations in challenging conditions, outperforming existing single-particle tracking algorithms. Finally, we show that traditional sptPALM experiments can be performed up to 5x faster without loss of information with current localization methods. This is especially useful in biological conditions where a short event needs to be investigated, such as recovery from DNA damage, or protein organisation during, e.g., phage infection or cell division. TARDIS is provided as an easy-to-use software tool via a GUI or implementable script-wise (https://github.com/kjamartens/TARDIS-public).

TARDIS is robust with respect to all causes that increase spt complexity: (1) high density of trajectories, blinking particles or short trajectories, (3) unstructured spurious localizations, and (4) structured spurious localizations. Additionally, we investigated scenarios that could hypothetically bias TARDIS and discuss these in Supplementary note 6. This investigation establishes that TARDIS bias is only introduced when the inter-particle linkage histogram (red in Figure 1) actually contains information on intra-particle linkages (i.e., TP trajectories). This can normally be circumvented by determining the τ at which intra-particle linkages are exclusively present via a statistical test (1-sided Wilcoxon, see Supplementary note 3). Only if the experimental particle bleach time (also taking, e.g., movement out of the FoV or focus into account) is on the same order of magnitude as the experimental time, no proper inter-particle linkage histogram can be created, and TARDIS will be biased.

Importantly, our investigations revealed that TARDIS cannot ‘fail silently’, i.e., provide data that at glance looks accurate, but contains errors: TARDIS failures due to wrong localization shows clear errors in the residuals of the fit (Supplementary note 6). A recent computational method that performs an analysis similar to our TARDIS-JD extraction, but with inter-particle data obtained from τ = 0 frames, further emphasizes the robustness of our approach^36^ (Extended data figure 10). We believe this high robustness of TARDIS is especially useful in spt(PALM) where the dynamics and localization are coupled, and a high local density influences the perceived dynamics.

TARDIS can be used on its own, or alternatively, the output of TARDIS can be used as accurate *a priori* information in tracking software, such as the Bayesian spt software *swift*. Via this combination (Figure 1b, 2c), accurate tracking is ensured while full trajectory information is extracted. We show that with the introduction of TARDIS, and especially combined with *swift*, the current major limitation for high-throughput spt(PALM) analysis is high-density localization of mobile particles. Current localization implementations specifically designed for high-density datasets focus on immobile particles, which introduces two artefacts when using on datasets of mobile particles: (1) motion-blurred localizations are often interpreted as multiple static particles, and (2) information on multiple frames is included, which collapses localizations of close-by particles on consecutive frames to the same location, rather than accurately deducing a small jump distance. Artefact (1) can be remedied via stroboscopic illumination^37^, but this cannot always be implemented due to hardware limitations and light dose considerations on live cell health. Artefact 2 shows the clear need for high-density localization algorithms optimized for mobile particles. Throughout this manuscript, we show that TARDIS is unaffected by densities of about an order of magnitude higher than can currently be accurately extracted from current state-of-the-art localization algorithms that can handle densities up to 5 loc/μm^2 13,14^.

In conclusion, TARDIS is a software platform that negates all traditionally avoided circumstances in spt(PALM), such as high density or strongly blinking particles. TARDIS therefore opens up the possibility of performing data analysis on mobile particles with paradigm-shifting conditions and enables the exploration of novel experimental designs.

## Acknowledgements

This work was financially supported by funding from a VLAG PhD-fellowship (J.H.), start-up funds at Carnegie Mellon University (B.T., K.J.A.M., and U.E.), the NSF AI Institute: Physics of the Future (NSF PHY-2020295) (U.E.), start-up funds at Bonn University (B.T., K.M., and U.E.), an Argelander Starter Kit at the university of Bonn (K.J.A.M.), and the Alexander von Humboldt Foundation (K.J.A.M.). We acknowledge the valuable input from group meetings from all members in the U.E. and J.H. laboratories.

## Data availability

All data underlying this study is available on doi:10.5281/zenodo.7900405

## Code availability

The custom TARDIS software used in this manuscript is provided as supplementary data and can be accessed on https://github.com/kjamartens/TARDIS-public.

## Author Contributions

Conceptualization: K.J.A.M. Data curation: K.J.A.M. Formal analysis: K.J.A.M. Funding acquisition: K.J.A.M., J.H, U.E. Investigation: K.J.A.M., B.T. Methodology: K.J.A.M., B.T., J.H, U.E., Project administration: K.J.A.M, J.H, U.E. Software: K.J.A.M. Supervision: K.J.A.M., J.H, U.E. Visualization: K.J.A.M. Writing–original draft: K.J.A.M. Writing–review and editing: All authors.

## Competing interests

The authors declare no competing financial interests.

## Extended Data figures

**Extended Data figure 1:**
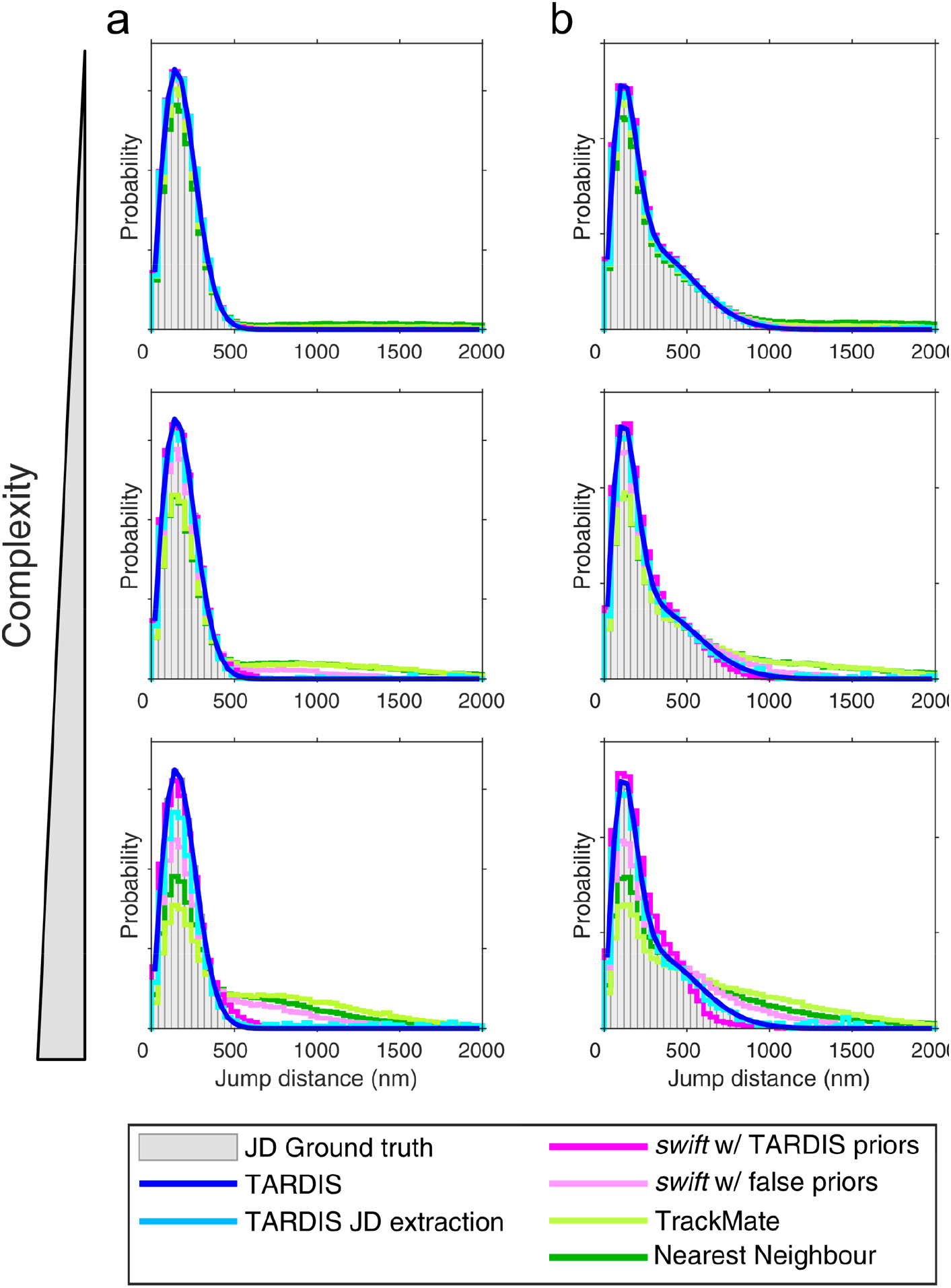
Performance of TARDIS and spt tracking methods for a single diffusive population and two diffusive populations at increasing complexity, visualised on a linear x-axis. Performance of TARDIS is compared to the Bayesian spt software swift, the LAP tracker in TrackMate^32,33^, and a simple nearest-neighbour analysis. This data is also presented in Figure 1b and Figure 2c.

**Extended Data figure 2:**
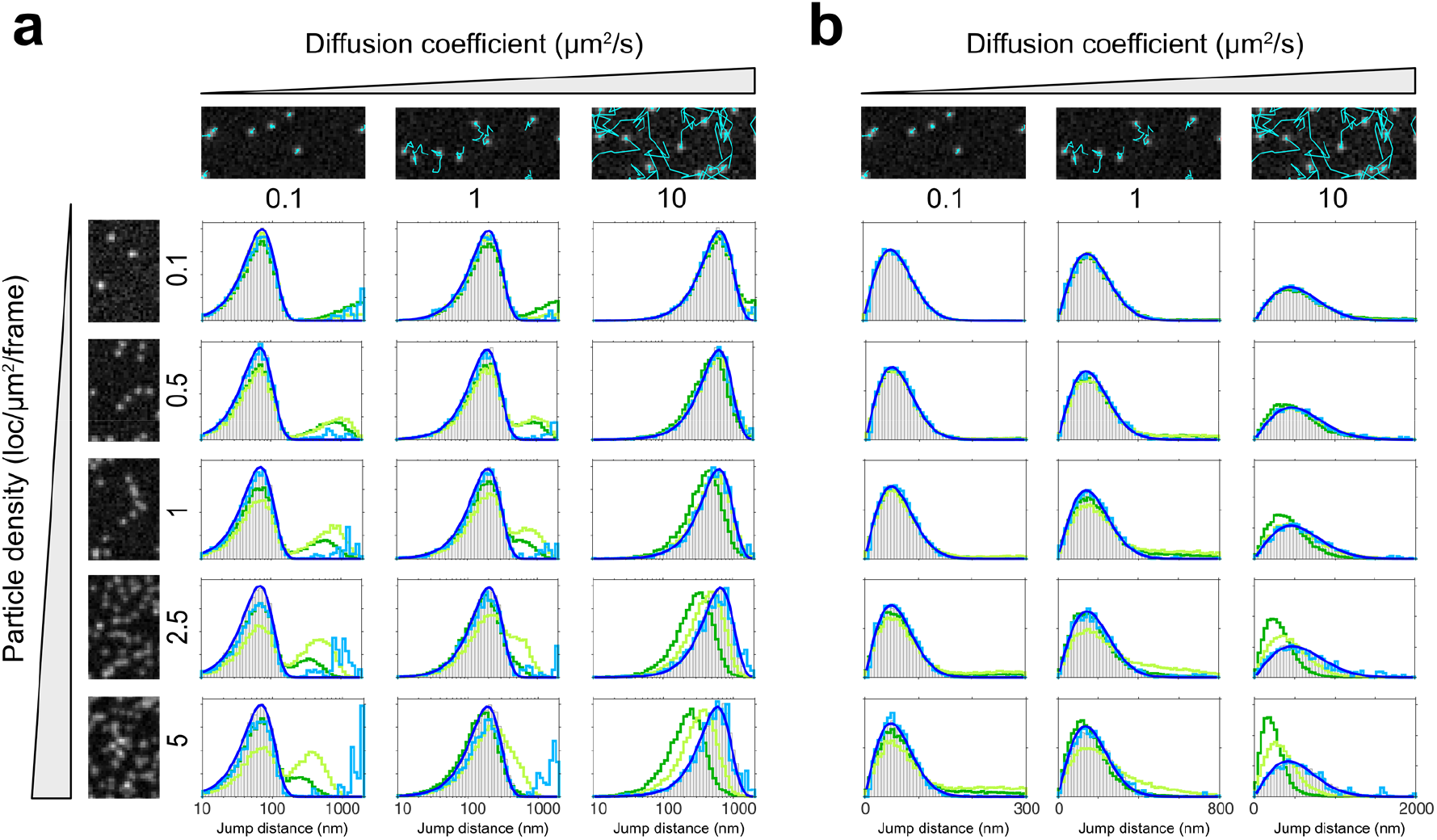
The full dataset as presented Figure 2a, analysed via TARDIS (blue), TARDIS-JD-extraction (light-blue), TrackMate-LAP ^32,33^ (light green) and nearest-neighbour tracking (dark green). (a) and (b) represent the same datasets, but visualised on a logarithmic (a) or linear (b) x-axis. Note the changing jump distance x-axis scaling in (b). The TARDIS fit data is also presented in Extended Data figure 3a.

**Extended Data figure 3:**
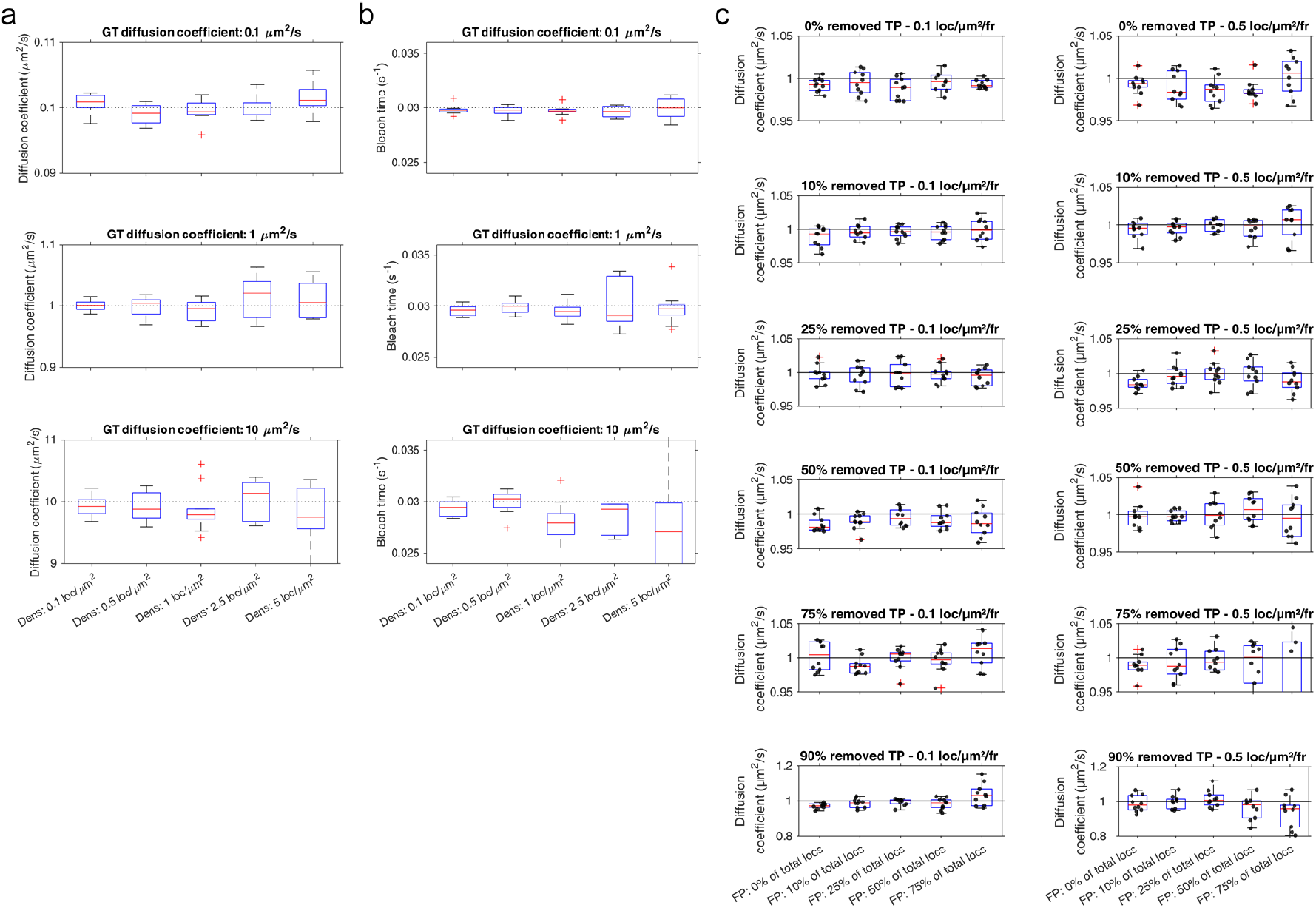
Detailed information on diffusivity and bleach time obtained from TARDIS fitting. (a) obtained diffusion of datasets presented in Figure 2a, and (b) obtained bleach time of datasets presented in Figure 2a, showing no bias in either over the complexity range. (c) Individual fit information of data presented in Figure 2b. For every condition, 10 repetitions were analysed with random start positions in TARDIS. The found diffusion coefficient is visualised (scatter points represent individual measurements). Note the changing y-axis at 90% removed true positives in (c). Abbreviations used: TP: True Positives, FP: False Positives, fr: frame, locs: localizations

**Extended Data figure 4:**
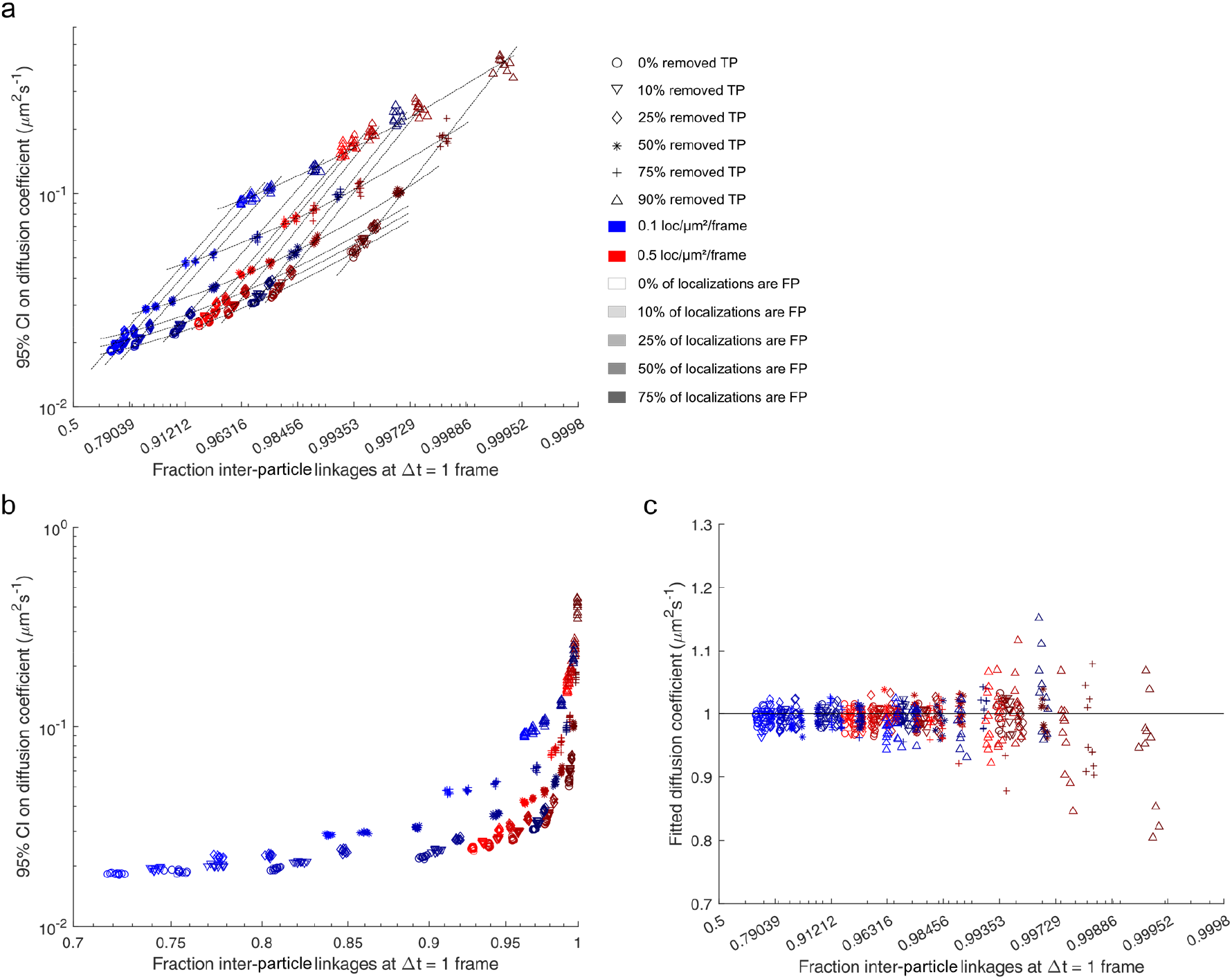
Effect of the fraction of inter-particle linkages on diffusion coefficient accuracy. Analysis of 95% confidence interval (a,b), and fitted diffusion coefficient (c) as a measure of the inter-particle fraction. Dotted lines in (a) are added for clarity. The underlying analysed data is the same as shown in Figure 2b. Reasons for increased fraction of inter-particle linkages are clarified via marker type (TP removal), marker colour (TP localization density), and marker darkness (FP introduction). Note that a and c have non-linear x-axis (a and b contain the same information, but with different x-axes).

**Extended Data figure 5:**
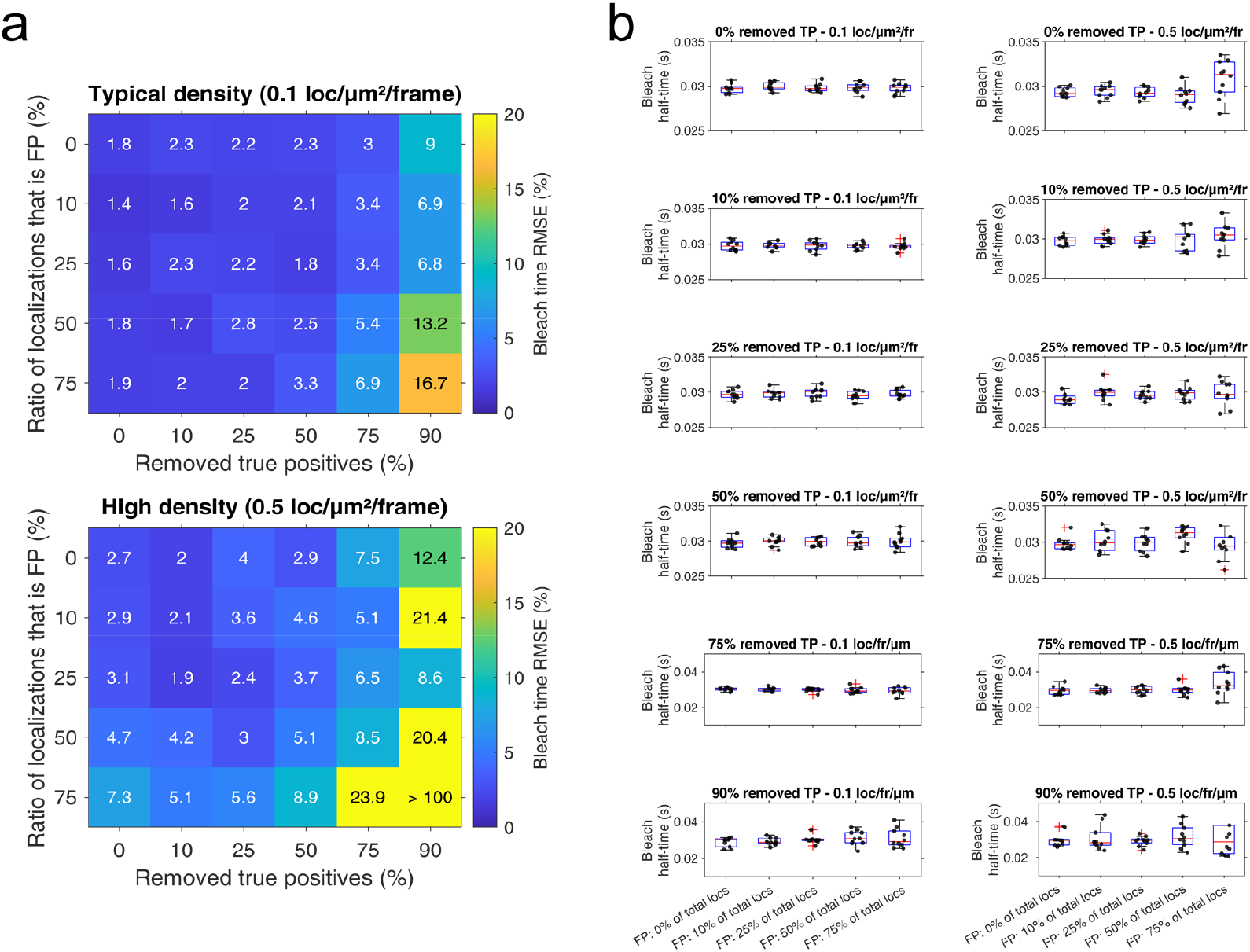
Detailed information on TARDIS fitting results of bleach time of single diffusive population with added noise and blinking chance. Fit information of bleach characteristics corresponding to the data presented in Figure 2b (main manuscript). For every condition, 10 repetitions were analysed with random start positions in TARDIS. (a) RMSE of the bleach half-time for all conditions (b) The found bleach half-time is visualised (scatter points represent individual measurements). Note the changing y-axis at 75% removed true positives. Abbreviations used: TP: True Positives, FP: False Positives, fr: frame, locs: localizations.

**Extended Data figure 6:**
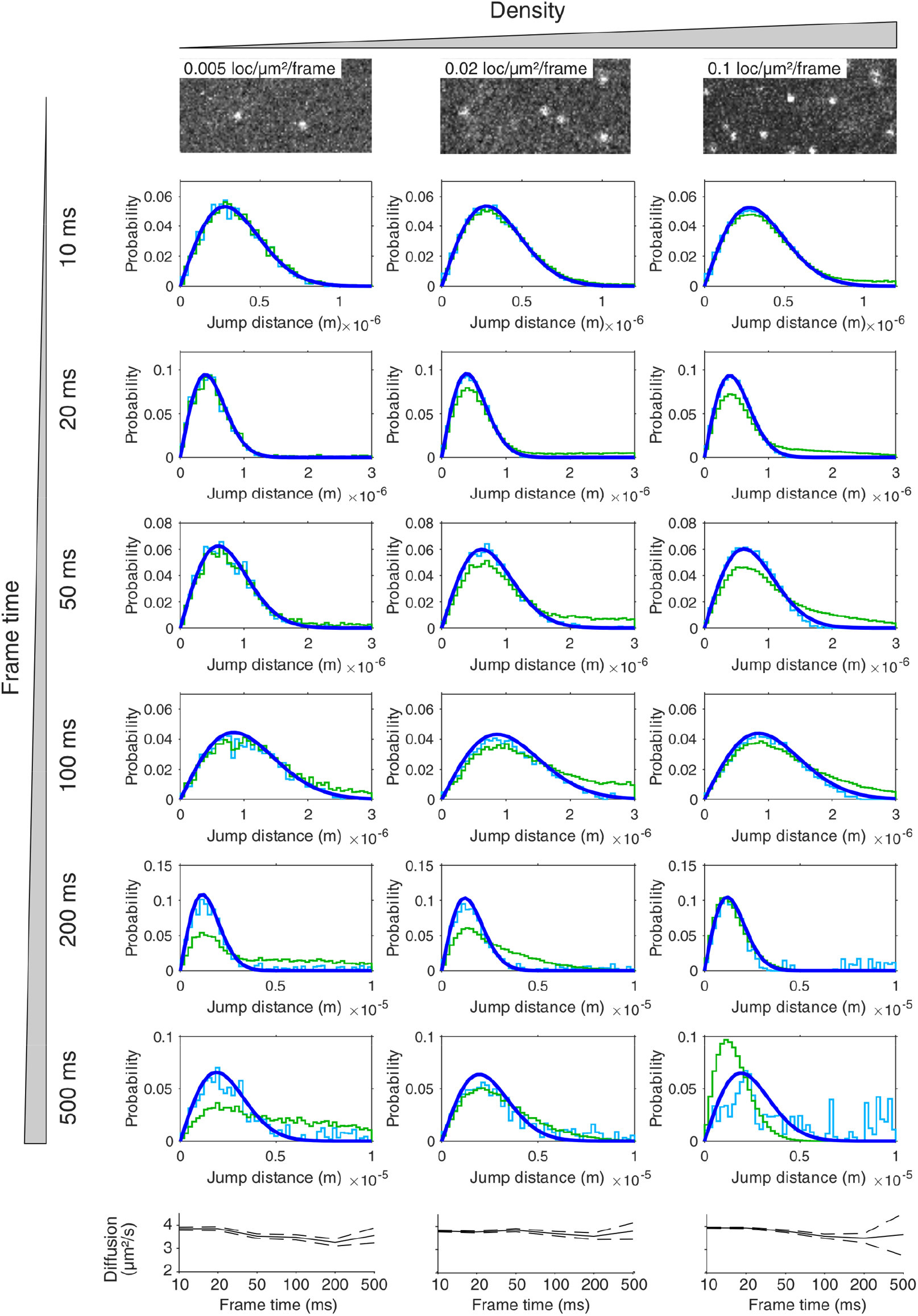
Diffusion analysis of fluorescent beads at varying frame times. The same information as presented in Figure 2d, but with additional frame times in between those shown in the main manuscript. The excitation time on every frame is kept constant. The small decrease in obtained diffusion coefficient as a function of frame time is explained by particles having a higher chance to move outside the field-of-view with larger jump distances.

**Extended Data figure 7:**
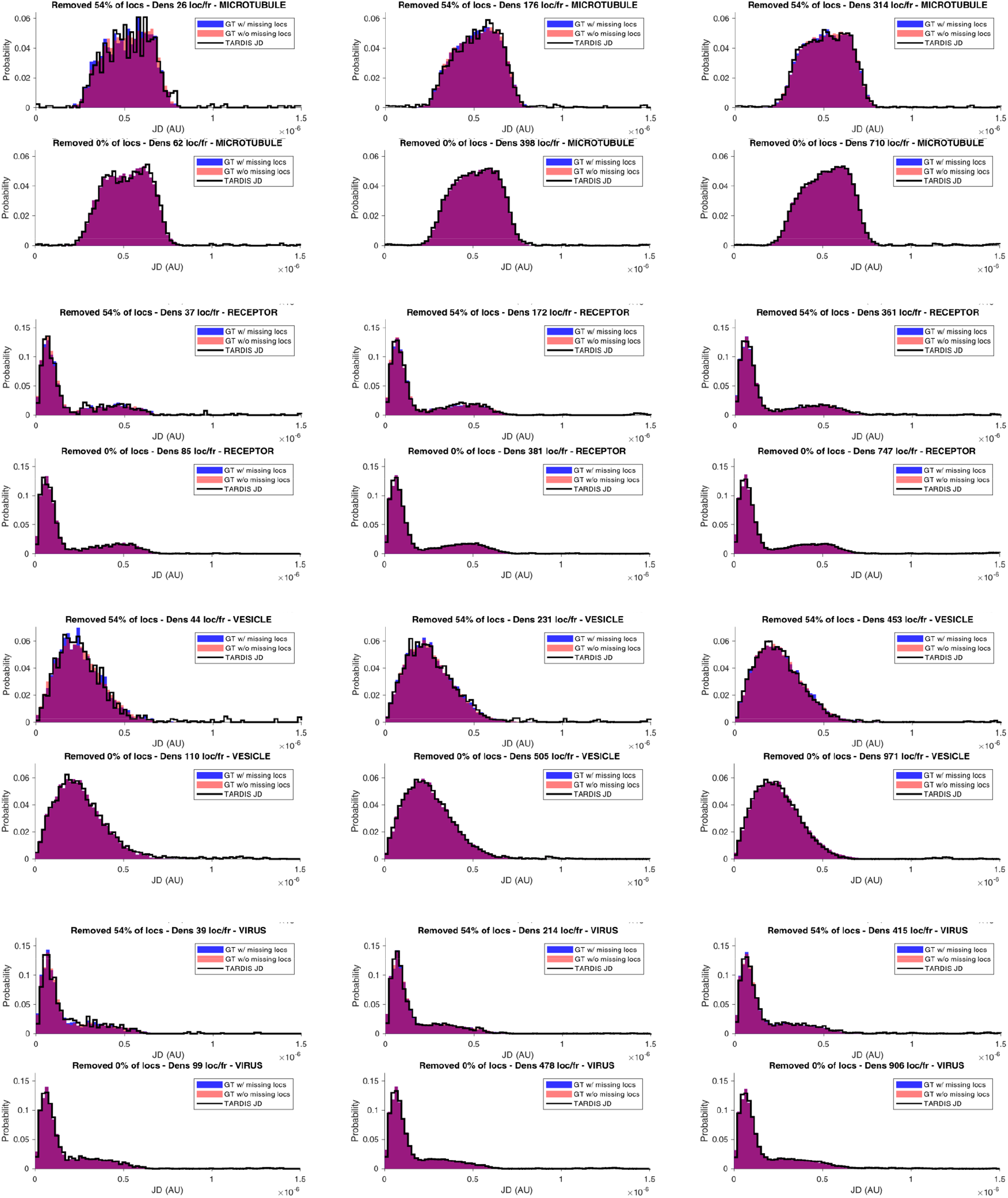
TARDIS-JD extraction from Chenouard et al. Tracking data from Chenouard et al^19^ has been deteriorated (removing 54% of localizations), analysed via the ‘extract JD’ -function of TARDIS, and compared to the ground-truth (GT) data. Four different conditions are analysed: MICROTUBULE, RECEPTOR, VESICLE, and VIRUS, corresponding to [constant velocity], [tethered motion, switching, any direction], [Brownian motion, any direction], and [same direction dynamics, switching between Brownian and linear] dynamics, respectively. Densities are indicated in subplot titles, while the field-of-view is ∼50-by-50 μAU in size. In all scenarios, TARDIS accurately extracts the ground-truth data, and the level of noise is decreasing with decreasing localization removal. The following TARDIS settings were used: Δt bins of 1-3; maximum jump distance of 1e-05 AU; background frames starting at frame-shift of 35, using in total 50 frames; 300 BG bins starting at 3.5e-06 AU.

**Extended Data figure 8:**
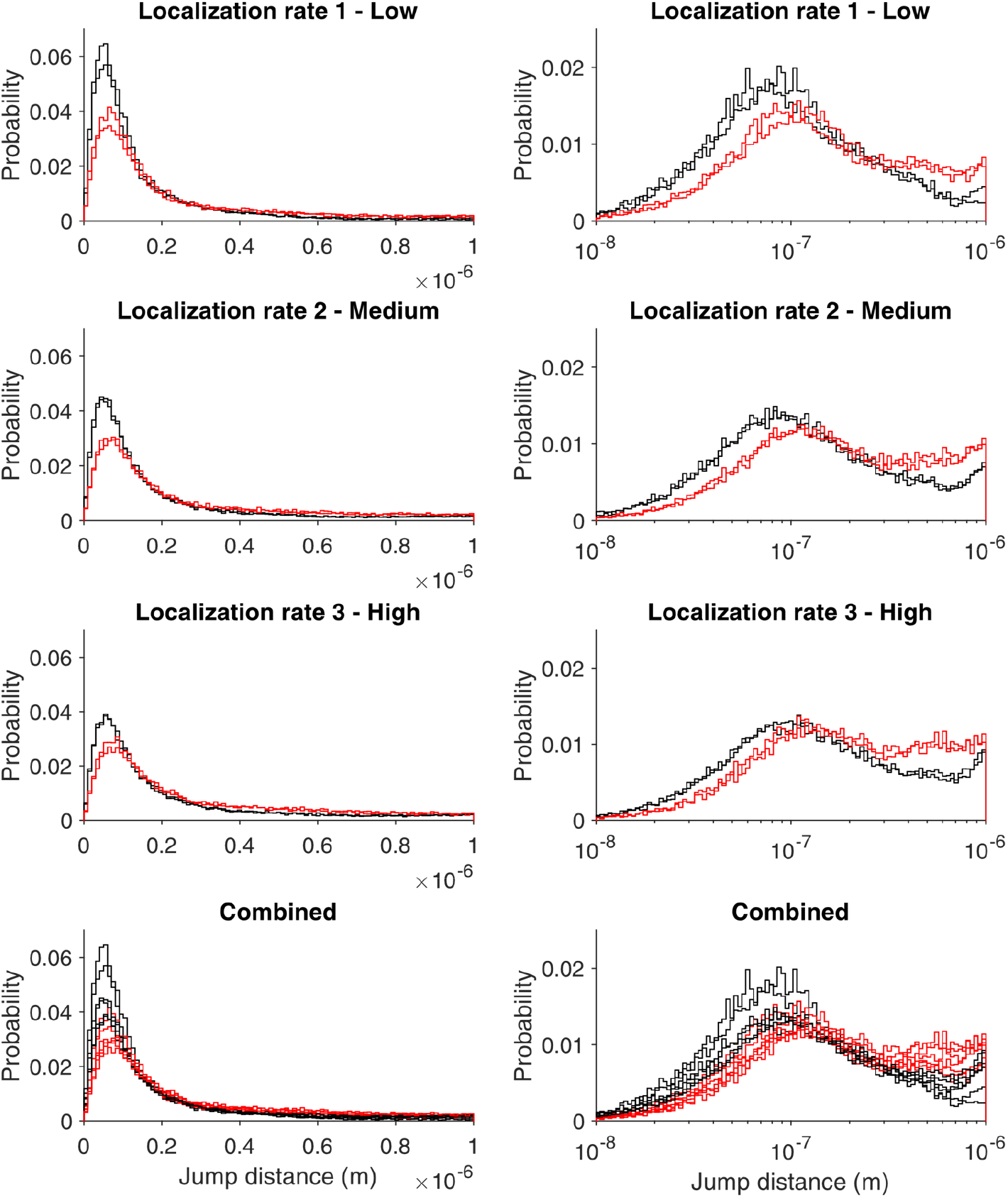
E. coli RNA polymerase jump distance analysis after nearest-neighbour tracking. Jump distance of the same data presented in Figure 3b, but tracked via nearest-neighbour tracking. Notice the changing peak position and abundance as a function of localization density. The data is shown in a linear X-scale (left) and logarithmic X-scale (right).

**Extended Data figure 9:**
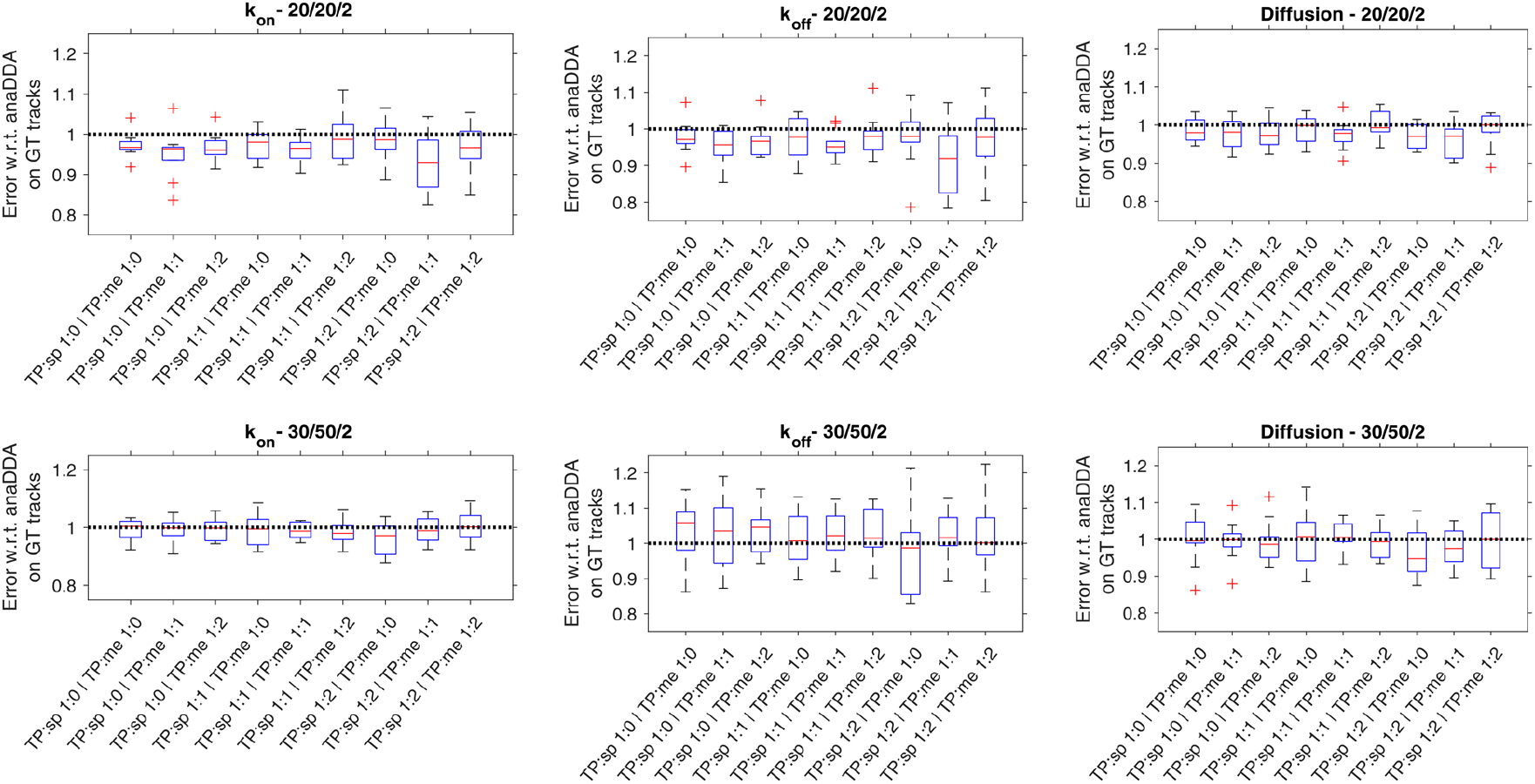
Detailed information on kinetically state-changing particles. Individual fit information of data presented in Figure 3c, along with individual fit information for differing binding/unbinding kinetics (titles indicate *k*_*on*_ / *k*_*off*_ / Diffusion coefficient). For every condition, 10 repetitions were analysed with random start positions in TARDIS, and compared to analysing the same data with anaDDA on the ground-truth trajectory data. TP, *Sp* and *me* indicate true positive, spurious, and membrane localizations, respectively.

**Extended Data figure 10:**
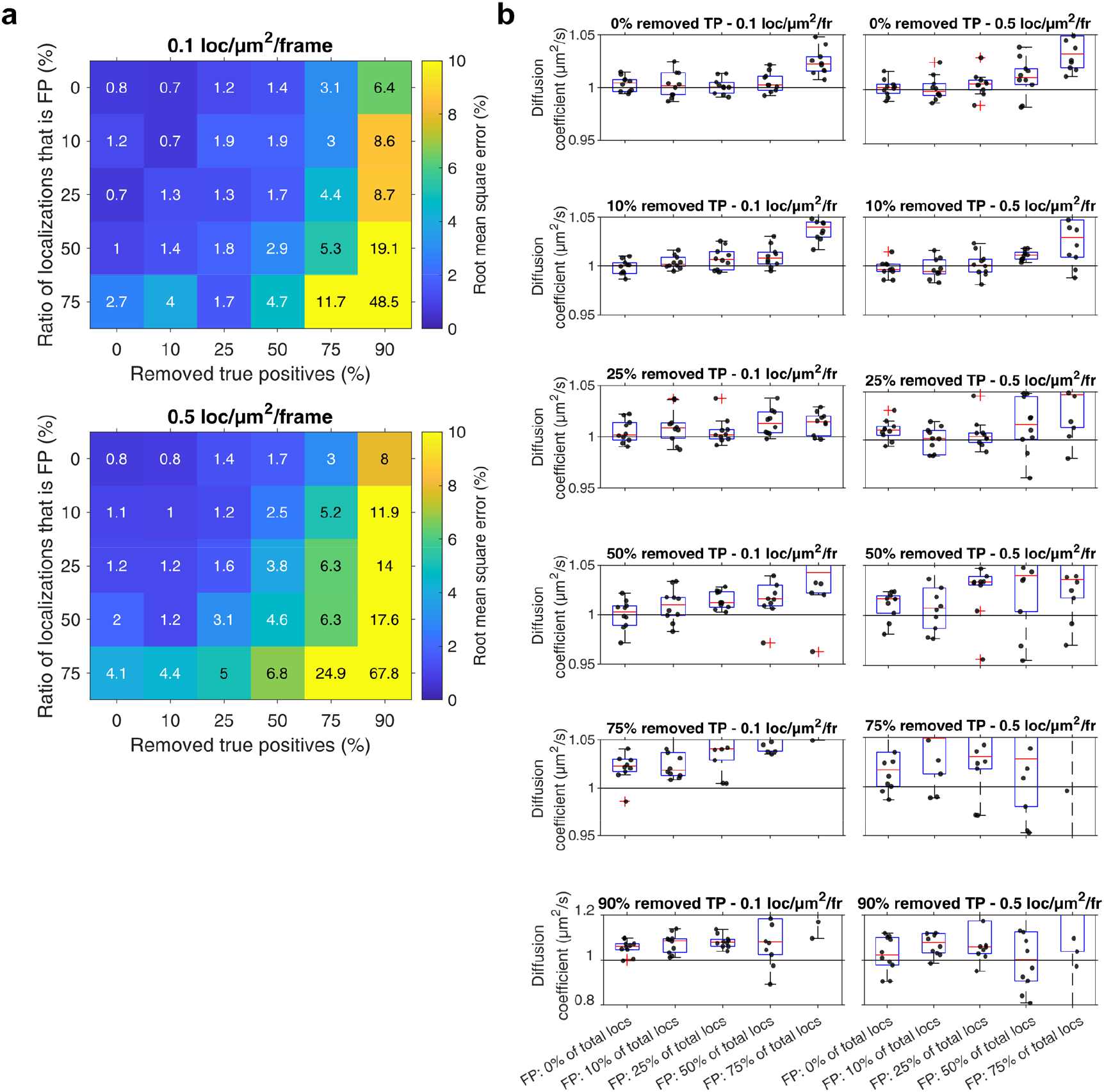
Accuracy of software by Wolf et al ^36^. The same data analysed by TARDIS in Figure 2b (a) and Extended Data figure 3b (b), analysed via the DANAE software^36^, which effectively only performs TARDIS-JD extraction. This is then fitted with a diffusive model afterwards. The accuracy, especially at high complexity scenarios, is worse compared to TARDIS. Additionally, DANAE shows bias towards too high values (right), which is caused by imperfect inter-particle distance distribution subtraction.

## Notes

https://zenodo.org/record/7900405

### Competing Interest Statement

The authors have declared no competing interest.

