## Supplementary Information for "Temporal analysis of relative distances (TARDIS) is a robust, parameter-free alternative to single-particle tracking"

### Supplementary Note 1: complete overview of computational aspects of TARDIS

This section contains an in-depth overview of the computational aspects of TARDIS. It starts with descriptions of the inter- and intra-particle distance distributions (red and blue lines in Figure 1a-b, respectively), then continues with a succinct description of the fitting procedure.

### Inter-particle distance distribution

The inter-particle distance distribution is a result of linking localizations that are not belonging to the same molecule (red lines in Figure 1b). Therefore, these linked distances are governed by the distribution of all particles, rather than by the diffusive behaviour of the particles. It is dependent on the size and shape of the region of interest, and on potential density distributions of particles, such as when particles are embedded in cells. Therefore, the inter-particle distance distribution cannot be described with a continuous analytical formula and can vary substantially between datasets[1,2]. A single inter-particle distance distribution is created from the complete localization dataset and stored for further use.

### Intra-particle distance distribution

The distance distribution obtained from intra-particle jump distances (blue lines in Figure 1b) is dependent on the investigated system. We are currently restricting to three scenarios, but TARDIS can feasibly be expanded to more scenarios, given that it can be well described by analysing jump distance distributions. Three scenarios are a single diffusing species with a given diffusion coefficient and localization accuracy (shown in Figure 1b); two diffusing species with different diffusion coefficient and fractions, but identical localization accuracy; and a single species which interchanges between two diffusive states (i.e. a low-diffusion and a high-diffusion state) with given $k_{slow\rightarrow fast}$, $k_{fast\rightarrow slow}$, $D_{fast}$, (and $D_{slow}$ = 0 μm$^2$/s), and localization accuracy, possibly additionally including a stable diffusing species with identical localization accuracy[3]. All these scenarios can be accurately described by analysing jump distances at varying time shifts.

### Fitting procedure

Obtained distance distributions, consisting of a combination of inter-particle and intra-particle distances, are fitted with a combination of inter-particle distance distribution (obtained from analysis at one or multiple high values of τ) and intra-particle distance distributions (dictated by an analytical formula). While a maximum-likelihood estimation (MLE) can be obtained for the analytical intra-particle distance distribution, this is not the case for the inter-particle distance distribution, as it is a discrete distribution. Therefore, we obtain the MLE of a datapoint belonging to the inter-particle distance distribution by interpolation between known positions at the inter-particle distance distribution.

Specifically, analysis with TARDIS is split in two parts. First, an estimation fit is performed to get approximate values for all important parameters to prevent local minima by the final fit. This estimation fit can be skipped by the user if requested. Next, the final fit is performed starting at the approximate values (but not restricted in any way).

First, the data for all τ bins is generated (i.e. as shown in Figure 1b). For this, all relative distances between localizations with a specific time shift are obtained from the dataset (i.e. for τ = 1, 2, 3, …), with a user-defined maximum jump distance to reduce complexity (but which can and should be substantially higher than the maximum expected jump distance). Additionally, a 'Background-JD' dataset is obtained, by obtaining these relative distances at one or multiple high τ intervals. When more τ intervals are used, more datapoints are gathered in the 'Background-JD' dataset, but this also increases the downstream analysis time.

Next, the 'Background-JD' and the various τ-interval datasets are histogrammed, and the data complexity is reduced by interpolating these histograms at ~300 points (user-definable).

***Estimation fit and TARDIS JD extraction***

With the TARDIS estimation fit, the ratio of data corresponding to the 'Background' rather than the actual data is approximated at every τ-interval by a root-mean-square-error (RMSE) calculation of the difference between the normalized τ-interval-histograms and the 'Background'-histograms multiplied by values between 0 and 1, only taking *x*-values over a certain user-defined threshold (normally ~75% of the maximum distance; e.g. 0.75 μm in Figure 1b) into account. It is expected that above this threshold, only background information is present, and no signal data is present. This effectively finds the ratio of background to signal where high *x*-values are fully explained by the 'Background'-data.

Next, curves are generated based on the interpolated τ-interval histograms, where the interpolated 'Background'-histogram is subtracted, based on the ratio of background to signal. This curve is used to create a set of 'fake' jump distance arrays, where 10.000 entries randomly populate this curve. Additionally, because the curves can have negative values, this is offset by a constant, so that the curves that are the basis for the 'fake' jump distance arrays are always positive. These curves can then be used to obtain a list of expected JDs (which is the output of the TARDIS JD approach). The length of the JD list is determined by the length of the τ-interval lists, corrected for the background-to-signal ratio.

For the TARDIS estimation fit, these 'fake' jump distance arrays are then fit with the distribution of choice alongside the constant offset. The specifics of this MLE-fit are explained below. This fit is globally performed over all τ-intervals.

***Full fit***

The full fit is started with the output parameters of the estimation fit. An MLE fit is instigated which contains the complete τ-interval datasets, and fits a combination of populations and 'Background'-data, based on the interpolated histograms previously obtained.

***MLE fit details***

For a single population fit, the likelihood that the input values belong to a curve is calculated with the following formula:

$$L = x \cdot e^{\frac{-x^2}{4\cdot\left(D+\frac{\sigma^2}{dt}+\frac{st}{6\cdot dt}\cdot D\right)\cdot dt}} + offset \qquad \text{Eq. 1}$$

in which *L* represents the likelihood, *x* the input data, *D* the diffusion coefficient corrected according to Berglund et al.[4], *σ* the localization precision, *dt* the time between the datapoints (i.e. the frametime multiplied by the time shift τ in frames), *st* the excitation pulse duration, and *offset* the linear offset introduced in the estimation fit, or 0 in the full fit. Note that this is calculated for all τ-intervals simultaneously by creating a single array with input data to be fitted, containing input data of all τ-intervals, and then changing *dt* in Eq. 1 to correspond to this.

This likelihood is then normalised via pseudo-integration. Briefly, the integral of the curve of either the signal or the background is estimated by taking 50.000 (but user-definable) bins of the Eq. 1, or the interpolated 'Background'-curve, respectively. The likelihood of all values is then normalized according to this value.

For a dual-population fit, Eq. 2 is used, and the further calculation is similar to the one described above.

$$L = \alpha \cdot \left( x \cdot e^{\frac{-x^2}{4\cdot\left(D_1+\frac{\sigma^2}{dt}+\frac{st}{6\cdot dt}\cdot D_1\right)\cdot dt}} \right) + (1-\alpha) \cdot \left( x \cdot e^{\frac{-x^2}{4\cdot\left(D_2+\frac{\sigma^2}{dt}+\frac{st}{6\cdot dt}\cdot D_2\right)\cdot dt}} \right) + offset \qquad \text{Eq. 2}$$

For anaDDA, a similar procedure is used, but rather than fitting jump distances, the apparent diffusion coefficient is fitted[5].

Finally, the total likelihood that the datapoints belong to the input parameters is calculated by adding the likelihoods of one or multiple populations together, normalized based by their ratio, and additionally adding the likelihood of belonging to the ‘Background’-curve, based on the background ratio.

### Supplementary note 2: Fitting the bleach kinetics

Since the trajectory length, and thus contribution of correct linkages to increasing τ values, is directly dependant on apparent (i.e. also influenced by factors such as moving out of focus/FoV) fluorophore bleach kinetics, the rate that the intra-particle population decreases can be fitted with this bleach value, if chosen.

Assuming typical bleach kinetics, a fluorophore trajectory length is governed by:

$$P = 2^{\frac{-1}{\lambda} \cdot t}$$

Where $P$ is probability, $\lambda$ is bleach half-time, and $t$ is trajectory length ($\lambda$ and $t$ in same units). Since a trajectory length of e.g. 4 frames contributes to 3x one-frame jumps, 2x two-frame jumps and 1x three-frame jumps, the contribution to TARDIS over all τ is given by:

$$P(\tau) = \sum_{n=1}^{\infty} \sum_{\tau=1}^{n-1} (n - \tau) \cdot 2^{-\frac{1}{\lambda}\tau}$$

Since there is no strict maximum trajectory length for natural bleach kinetics, this should in theory be computed for n = 1 → ∞. The mild assumption of calculating this to n = ~500 gives the same result. This probability is further normalized and corrected for the total number of jump distances found at all values of τ in TARDIS.

### Supplementary note 3: Estimation of longest trajectory length

The estimation for the longest trajectory length is not trivial. In many spt experiments, the bleach time of the used fluorophores is governed by an exponential decay. This in turn indicates that with the number of fluorophores increasing towards infinity, the maximum bleach time in the experiment also increases towards infinity. In addition, the trajectory lengths normally obtained from traditional spt approaches do not take blinking into account (i.e. these are underestimating the true trajectory length), meaning that trajectory lengths quantified that way are not suitable for use with TARDIS.

A way to autonomously estimate the longest trajectory length in the data, using only the data itself is described here. The basic principle is as follows: First, it can reasonably be assumed that at very long time shifts, the influence of unbleached fluorophores on TARDIS results is neglectable. For this, TARDIS uses a value of 25% of the total movie length. Then, the relative distance histogram obtained at these long time shifts can be obtained as described above. More relative distance histograms are then obtained at regularly (logarithmically) spaced positions from very short time shifts (e.g. 10 frames) to the very long time shifts. Finally, we can test whether the relative distance histogram obtained at long frame-delays is statistically different from all relative distance histograms obtained at other frame delays (Supplementary figure 1). This is done using a 1-sided Wilcoxon rank sum test (also known as Mann-Whitney U test), which tests whether the median of the histogram obtained from short time shifts is shifted towards lower values compared to the histogram obtained from the long time shifts (the histogram obtained from short time shifts is randomly sampled to obtain the same number of distances as the long time shifts histogram). If this median is indeed shifted, there is still a substantial amount of correlated trajectory data present. The first time shifts at which the median is no longer left-shifted (with $p < 0.01$) can be considered the lowest time shift that can be used as the longest trajectory length.

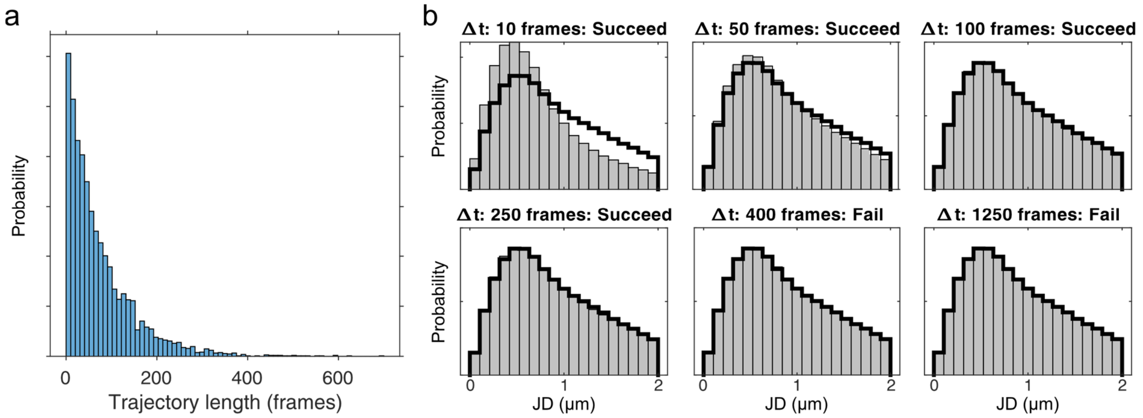


***Supplementary figure 1: Autonomous longest trajectory length estimation using the 1-sided Wilcoxon rank sum test.*** 5000 trajectories with a bleach half-time of 0.5 seconds (at 10 ms frametime) are simulated. (a) the ground-truth distribution of trajectory lengths. (b) relative distance histograms obtained at 10, 20, 50, 100, 250, and 1250 frame time shifts. These histograms are all tested with the 1-sided Wilcoxon test against the relative distance histogram obtained at 1250 frame time shift, and the Wilcoxon test (p<0.01) result is indicated in the subpanel. These results indicate that a 400 frame time shifts (but not a 250 frame time shifts) can reasonably be used as maximum trajectory length without biasing TARDIS results. In the test implemented in TARDIS, more than 6 time shifts are tested for.

### Supplementary note 4: Estimation of *swift a priori* parameters from TARDIS

*swift*, being a Bayesian-based method, requires *a priori* estimates for the parameters in its model. It is beneficial to provide accurate estimates to improve the accuracy of *swift*. The main parameters for *swift* are expected displacement, expected noise rate, and probabilities of bleaching, blinking, and reappearing of particles. TARDIS can be used to provide first estimates for *swift*, using the following logic.

A first consideration is that ideally, these parameters can be extracted by TARDIS without performing an analytical fit (i.e. by JD-extraction). The expected displacement can simply be estimated by providing the median jump distance of TARDIS JD extraction (Figure 1).

Obtaining the expected noise rate, probability of bleaching, and probability of blinking, however, is a more complex issue. As mentioned in Supplementary Note 1 above, all three of these factors contribute to the ratio between inter-particle linkages and intra-particle linkages. This entails that they cannot be individually extracted via TARDIS, except for bleaching rate, which can be extracted from the decrease of intra-particle linkage size over time-bins (see Supplementary Note 2). As the expected noise rate can be reasonably deduced via proper negative control experiments, we created a method to extract the blinking probability if an expected noise rate is provided.

The following analysis is used (Supplementary figure 2): The number of linkages between localizations per frame scales with true localization density (TP-dens), noise density (noise-dens), and is inversely proportional to blinking ratio. First, the total number of linkages between localizations per frame is calculated within the search radius of TARDIS as described above (i.e. logical additional output from analysis described in Figure 1b), and this is done over many time shift bins. Next, the number of linkages between localizations per frame only at high-τ values is determined (i.e. absolute number of linkages in Figure 1b at τ=3, but generally at high-τ values). This number should stay stable, but slowly decreases because τ becomes a more relevant fraction of total frame times at high τ. From this slow decrease, the slope and intersection of the y-axis is determined. Then, the difference between number of linkages at every frame between the high-τ-values and low-τ-values is determined at τ=1. Finally, all of this is normalized for TP density.

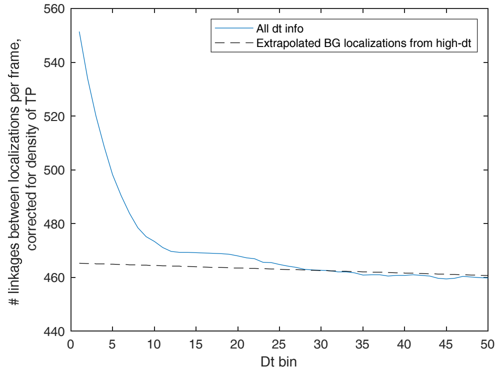


***Supplementary figure 2: Assessment of number of linkages between localization per frame***

The analysis described above is shown visually. The number of linkages between localizations per frame, corrected for the true localization density, is plotted as a blue curve. The number of linkages belonging to background localizations is plotted as a dashed black line. The difference at τ=1 is a measure for the number of linkages belonging to moving particles.

Next, separately from the analysis above, the number of linkages expected if no blinking is happening, is deduced as follows: From the total number of localizations in the dataset, the number of TP localization is calculated via the ratio of TP-dens to noise-dens. With the TARDIS-obtained bleach time (Supplementary Note 2, which is independent of blinking), the expected number of TP trajectories is determined from the number of TP localizations. By then assuming that the trajectories are evenly spread over all frames in the dataset, the number of linkages between the trajectories can be calculated, where it is assumed that no blinking is occurring. This is finally also normalized for TP-dens.

The ratio between these two analyses is a direct measure for the blinking occurring in the dataset, and is extracted like this. This analysis is used on the dataset described in Figure 2b to accurately obtain the blink fraction, especially accurately at low-to-medium complexity (Supplementary figure 3).

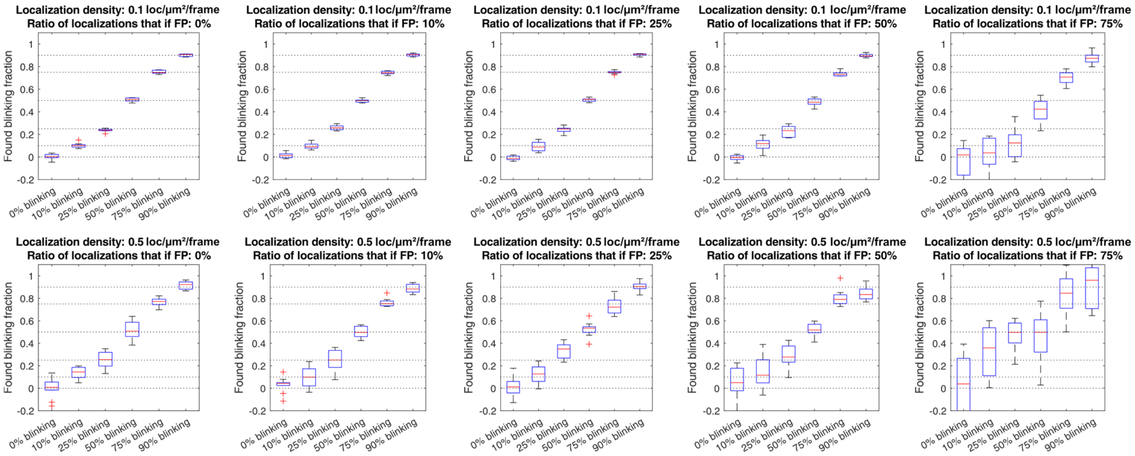


***Supplementary figure 3: Approximation of blinking fraction from TARDIS output.***

The data described in Figure 2b is used for this analysis. The only required input parameter is approximate noise density, which has been determined separately for every density/FP_ratio combination.

### Supplementary note 5: Complexity in single-particle tracking

We note that complexity for spt analysis can come from different sources, but have qualitatively similar effects on spt analysis. We briefly note these here as rationale for not elaborately testing across all possible complexity spaces.

- Increasing particle density versus increasing diffusion coefficient: in both cases, there will be an increase in the number of 'false particles' within the 'true search space' of 'true particles'. For example, an particle that moves on average 100 nm requires e.g. a search area of $3 \cdot \pi \cdot 0.1^2 = 94$ $\mu m^2$ (assuming a circular search space of three times the average jump distance). Complexity is increased by increasing the number of false particles in this search space, which can be increased by either increasing particle density or by increasing the search space (i.e. increasing the diffusion coefficient or increasing the frame time whilst keeping the excitation time constant).
- Increasing false positive localizations: Following the argument from above, increasing the number of false positive localizations directly influences the number of false particles in the search space, similarly complicating spt analysis.
- Increasing blinking fluorophores: Following the previous argumentation, by decreasing the average number of true particles in the search space, but keeping the false particles stable, the spt analysis is similarly complicated.

These sources of complexity do not contribute equally to spt analysis error rate, but in general can all be interchanged for one another with similar results. This is also shown in Extended Data figure 4.

### Supplementary note 6: Possible limits of TARDIS

In this paragraph, we investigated and reported on possible limits of TARDIS. We created custom Monte-Carlo simulations of different scenarios, and analysed the process and/or output of TARDIS to investigate their effect.

#### Effect of inaccurate localization at high localization densities

At high localization densities, not all underlying point-spread functions (PSFs) of different particles are fully spatially separated. Having overlapping PSFs may lead to wrong localization, most notably localizing two closely-spaced PSFs as a single one, with the found position (as a first approximation) being an average of these PSFs. If the positions of PSFs are randomly spaced, or structurally spaced without bias (e.g. in dSTORM microscopy), this creates unbiased errors, but no spatially biased errors.

In single-particle tracking, however, it is more likely that the position of the trajectory's particle in the next frame is closer by than (1) other trajectories, or (2) spurious noise. This is illustrated when comparing the TARDIS probability (blue) and inter-particle distance (red) in e.g. Figure 1b. This bias results in a shift of jump distances that are attributed as intra-particle towards higher values. Since this bias is strongest at low frame-delays and fully disappears at time shifts larger than the longest trajectory length, it effectively creates a correlation that only exists at high time shifts (i.e. when determining the inter-particle distance), but not at low time shifts. This concept is illustrated in Supplementary figure 4, where a low-density (0.1 loc/μm$^2$/frame) and higher-density (0.5 loc/μm$^2$/frame) dataset of a single-population, non-deteriorated dataset (i.e. similar to Figure 1c, lowest complexity, only increasing density) is analysed with TARDIS with three different localization methods: (1) raw input localizations, (2) localization of simulated PSFs from the raw input localizations with a multi-particle fitting routine (in ThunderSTORM), and (3) localization of simulated PSFs from the raw input localizations with a single-particle fitting routine.

We note that these effects will have similar consequences to all tracking methods (Supplementary figure 5), since the localizations themselves are deteriorated, rather than the analysis of them. To the best of our knowledge, no tracking methodology accounts for these effects. Finally, we assume throughout the manuscript that analysis is perfect accurately, and that the effects investigated here can be discarded.

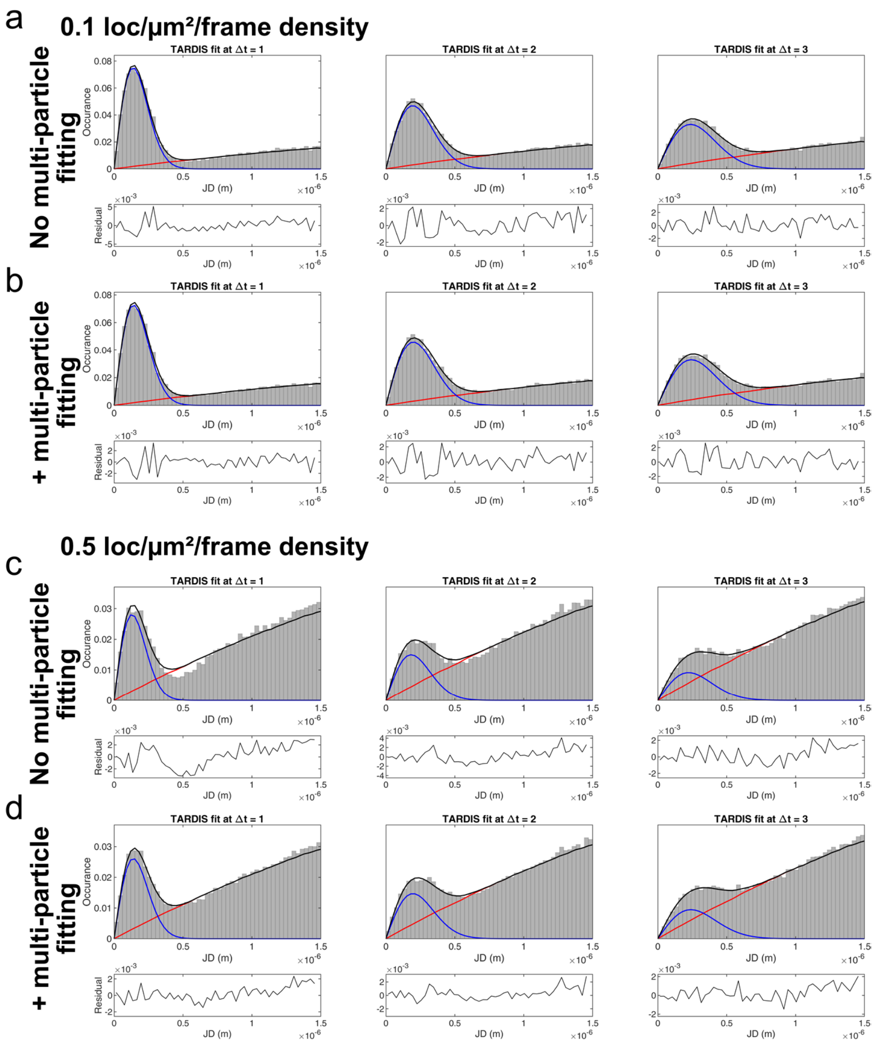


***Supplementary figure 4: Effect of multi-particle fitting artefacts on TARDIS***

Effect of multi-particle fitting and higher densities on TARDIS fit. A simulated dataset was created as a TIF movie (assuming PSFs are 2D Gaussians with randomly 1000-1500 photons on a 150 photon, Poisson-noised background, with sizes equivalent to 500-700 nm wavelength, using 100 nm pixel size), and was analysed with ThunderSTORM (1.5-2.0 pixel difference-of-Gaussian filter, local maximum at std(Wave.F1), fitted via 9x9-pixel Gauss-MLE fitting – where indicated, multi-particle fitting was used with maximum 6 fits, 1e-50 p-value, 1000-1500 intensity range. Found localizations below 800 photons, and found localizations that were on the same frame and less than 2*localization precision apart were discarded). (a,c): typical- and high- particle density analysed without multi-particle fitting present. (b,d): typical- and high- particle density analysed with multi-particle fitting. Note the gap at 500 nm found primarily in the 1 frame-delay image in (c).

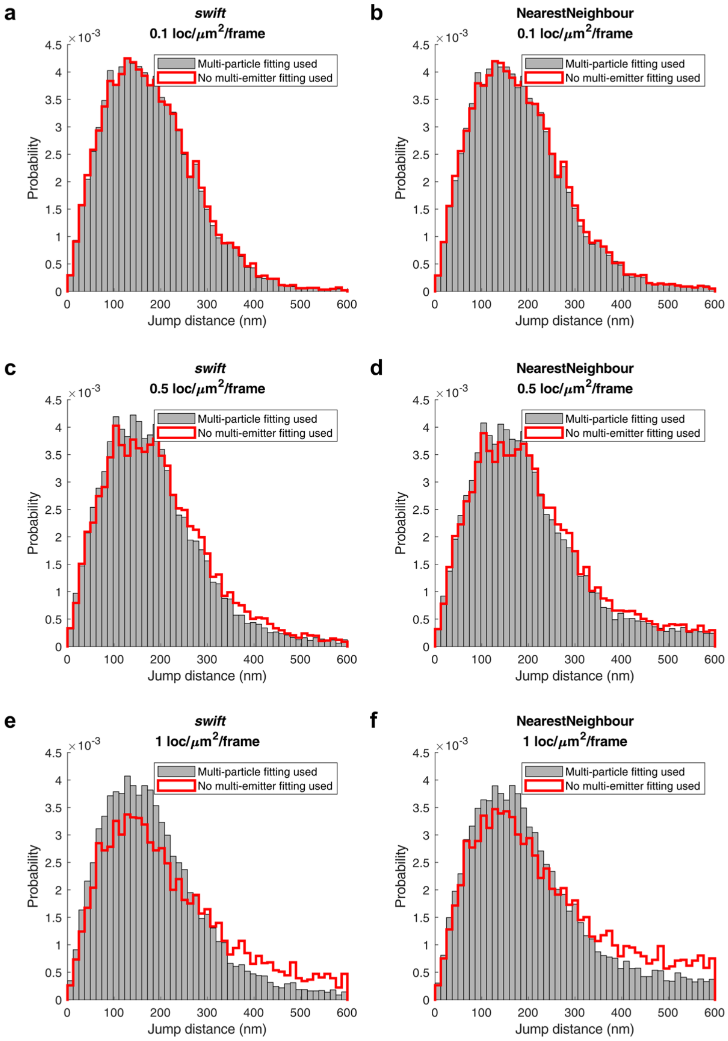


***Supplementary figure 5: Effect of multi-particle fitting artefacts on traditional tracking methods***

Effect of multi-particle fitting on simulated data at low (a,b), high (c,d), and very high (e,f) density of particles, analysed via *swift* (a,c,e) and nearest-neighbour tracking (b,d,f).

5

### Spatial heterogeneity combining density and diffusion

It can be envisioned that there is a correlation between location, fluorophore density, and fluorophore diffusion. We investigated if this has an effect on TARDIS by simulating a dataset (Supplementary Figure 6a) in which two populations exist at different sub-spaces in the dataset, either with D = 5 μm$^2$/s which start at maximum 1 μm distance from the center of the dataset (70% of trajectories), or with D = 1 μm$^2$/s, which start at a distance between 1 and 4 μm from the center of the dataset (30% of trajectories). 15 nm localization precision is used. We observed no difficulty for TARDIS to fit this dataset, with the obtained results being the following (Supplementary Figure 6b): $D_1$=0.94±0.091 μm$^2$/s at fraction 0.28±0.032, $D_2$=4.88±0.310 μm$^2$/s at fraction 0.72±0.032.

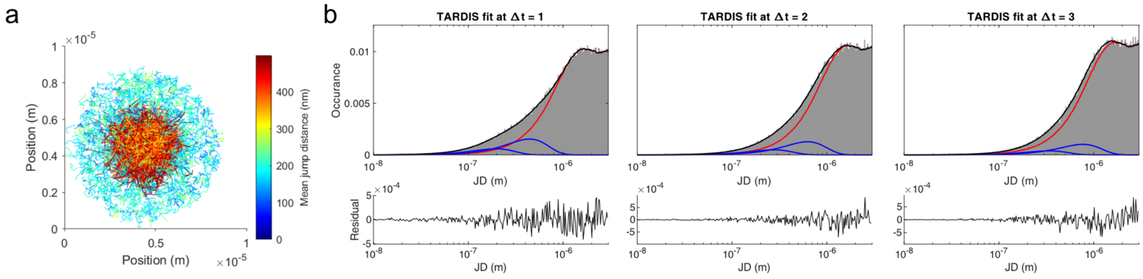


***Supplementary figure 6: TARDIS when applied to datasets with spatial heterogeneity in density and diffusion.***
(a) Overview of the simulated trajectories, colour coded by mean jump distance (b) TARDIS results of analysing the dataset.

### Localizations with high spurious blinking particles with similar size as the jump distance

We were wondering if having a small area (on the same spatial scale as the (mean) jump distance of diffusing particles) with high density of spurious localizations would have an effect on TARDIS. One could propose that there will be ambiguity in the TARDIS fit, since the inter-particle curve (red) overlaps with a population-like distribution at similar spatial scales compared to the intra-particle curve (blue). We simulated (Supplementary Figure 7a) a dataset like this as follows: a 1 μm$^2$/s diffusing population (black, trajectory length dictated by 2.2 frame half-time, start localization randomly in a 10x10 μm FoV, 30 nm localization precision, 10 ms frametime, 1000 frames, 10.000 localizations) was deteriorated with spurious localizations (red, 1000 localizations), and with highly concentrated and localized spurious localizations (blue, 2500 localizations). TARDIS analysis (Supplementary Figure 7b) shows that the spurious particle distribution is well-described by the intra-particle curve, and fits the inter-particle curve well, resulting in D = 0.99±0.036 μm$^2$/s.

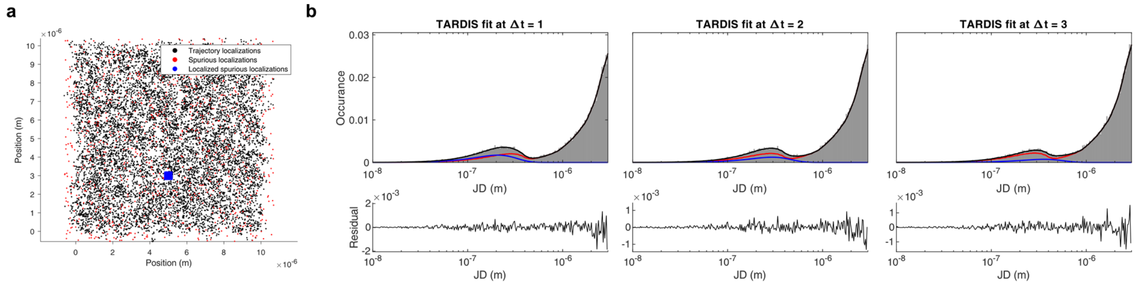


***Supplementary figure 7: TARDIS when applied to datasets with a localized area of high spurious blinking particles.***
(a) Overview of the simulated localization colour-coded as described in the text. (b) TARDIS results of analysing the dataset. Note that the inter-particle linkage distribution (red) clearly shows the effects of the localized spurious localizations.

### Non-bleaching particles

If particles are never bleaching, there can never exist an inter-particle only population (Figure 1). We investigate this effect on TARDIS fits by simulating the following scenarios:

In a single cell, 5 particles are simulated for 10.000 frames (0.01s frame-time) with state-changing kinetics of 20 $s^{-1}$ $k_{on}$ and $k_{off}$ ($D_{free}$ = 2 μm$^2$/s) which never bleach. This is conceptually similar to un-blinking, un-bleaching particles bound to protein of interests in a single prokaryotic cell. The resulting fit (Supplementary figure 8a) underestimates the diffusion coefficient, and the fit is visually imperfect (i.e. intra-particle curve does not perfectly describe the high-D*-population, and residuals show a sinusoidal pattern).

Next, we simulated 50 particles in a 2-dimensional plane (1.000 frames, 0.01s frame time, 2 um$^2$/s) that can move out of the field of view, but do never bleach. This is conceptually similar to un-blinking, un-bleaching fluorescent beads that freely diffuse. The particles start on a random frame in the movie, and the position at frames > 1.000 are used in the first frames of the movie instead. This sole condition (i.e. ability to diffuse out of the FoV) already satisfies TARDIS' requirement of finding an inter-particle only population, as can be see in the good resulting TARDIS fit (Supplementary figure 8b).

It should be noted that at these conditions, given there is enough data present, the inter-particle population (red) does still contain information of the diffusing species, which can be seen when simulating 500 particles with the same conditions, and investigating the residuals of the fits (Supplementary figure 8c). These indicate that the 'inter-particle' only population (τ = 500 frames and higher) still contains intra-particle linkages at high JD values (~1.5 μm), that are not found at dt = 1 to 3. However, this contribution is low, considering most of the diffusing species have diffused out of the field of view, and do not impact the absolute fitting values (2.00 +- 0.04 μm$^2$/s and 2.04 +- 0.02 μm$^2$/s, respectively).

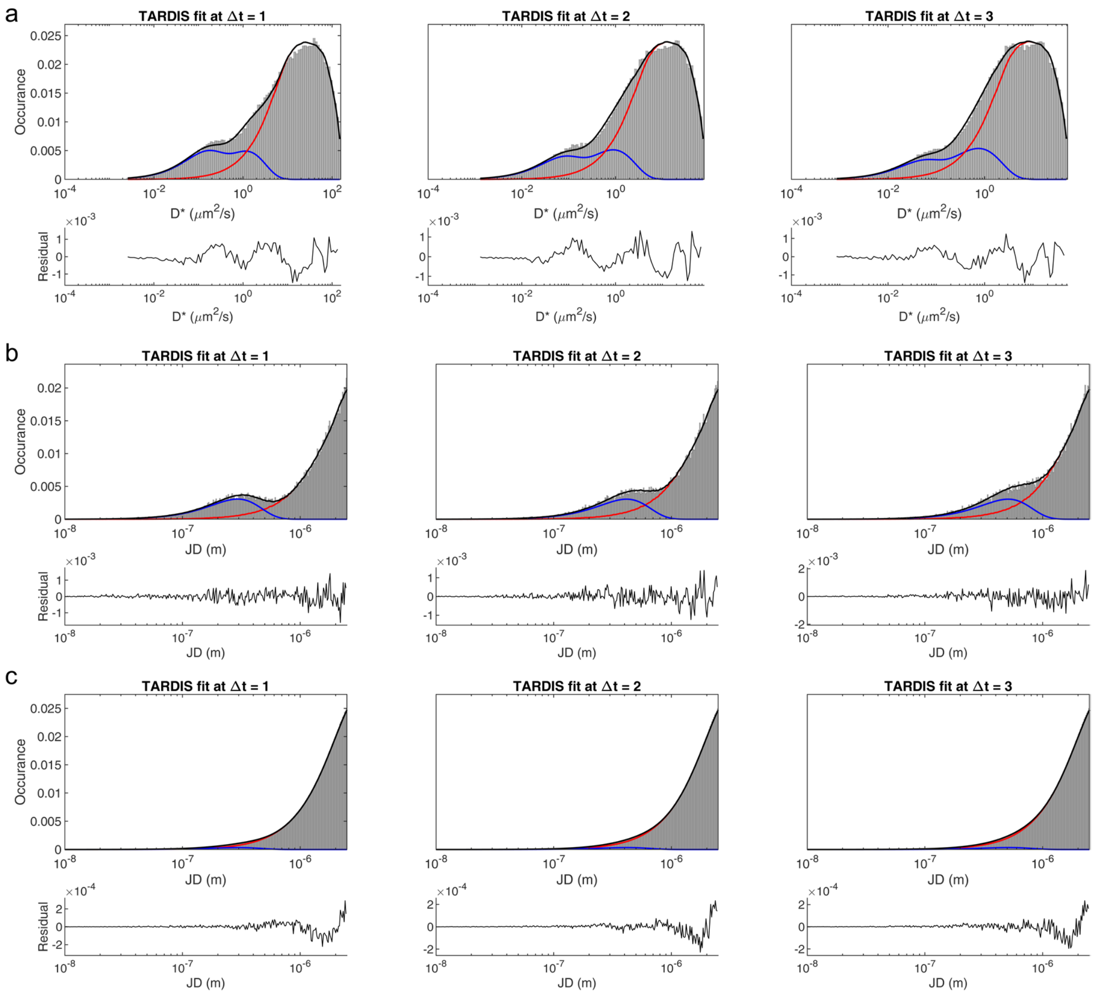


***Supplementary figure 8: Effect of non-bleaching particles on TARDIS fit***

TARDIS was used on datasets that do not have bleaching particles. (a) Imperfect fitting of 5 particles that move inside individual microbiological cells for 10.000 frames. (b) Perfect fitting of 50 particles that never bleach (1.000 frames long movie), that are allowed to move out of the field-of-view. (c) Same conditions as in (b), but with 500 particles, showcasing the small sinusoidal imperfection in the residuals of the fit. More detail about the simulations in the text above.
